# Contextual encoding of states and indices for episodic memories make goal-directed action selection flexible, transparent, and efficient

**DOI:** 10.1101/2025.10.25.684594

**Authors:** Yukun Yang, Wolfgang Maass

**Author notes:** Contributing authors.

## Abstract

When we want to solve a problem, our brain helps us by recalling related experiences. Furthermore, it can selectively recall only those experiences that fit especially well to the current context and contingencies. In addition, it provides explanations for chosen actions in terms of prior experiences. We show that experimental data on underlying neural codes of the brain, in particular contextual encoding of states and index cells for episodic memories, enable us to port essential aspects of the problem solving strategy of the brain into a simple algorithm. It only requires local online learning in shallow neural networks with low resolution of synaptic weights in order to explore a new environment, and it supports low-latency decision making. Hence it is especially suited for implementation in edge devices. Surprisingly, the performance of this light-weight algorithm on a standard benchmark task, navigation in Markov Decision Processes, is competitive with that of commonly used but computationally more demanding, less flexible, and less transparent reinforcement learning approaches.

## 1 Introduction

Whereas most machine learning methods store learnt knowledge in parameter values, there also exists a long tradition of exemplar-based methods in machine learning. Predictions are made there by comparing the current challenge with explicitly stored training instances (exemplars). The k-Nearest Neighbor (k-NN) algorithm and Support Vector Machines are classical examples for solving classification problems. Exemplar-based methods have also been integrated with DNNs (Deep Neural Networks) and LLMs (large Language Models), e.g. in the form of Neural Turing Machines or Retrieval Augmented Generation (RAG) for LLMs. If the goal is to learn behaviors instead of classifications, exemplars take the form of episodic memories (EMs) of past experiences. They play an important role in human strategies for solving Reinforcement Learning (RL) problems (Bornstein et al., 2017; Gershman and Daw, 2017; Bornstein and Norman, 2017; Biderman et al., 2020; Nicholas and Mattar, 2026). Case-based reasoning (Yan and Cheng, 2024) is an approach that ports some aspects of this solution strategy of the brain into AI-algorithms. However, it is less suited for navigation in random graphs or Markov Decision Processes (MDPs).

Several methods have already been proposed for integrating EMs into RL algorithms. They demonstrate some but not all of the functional goals that we want to achieve with our approach: Storing experiences in a goal-invariant way that enables navigation to arbitrary goals without relearning, low latency action selection that can be adapted online to new constraints or contingencies, explainability of action selection in terms of concrete prior experiences, online learning instead of offline training, and sufficiency of shallow neural networks with local synaptic plasticity that facilitates implementation in energy-efficient edge devices.

The benefits of EMs for guiding action selection in RL tasks was apparently first demonstrated by (Lengyel and Dayan, 2007). They showed that EMs support navigation to a fixed goal in tree-structured MDPs. The model-free episodic control approach of (Blundell et al., 2016) is more broadly applicable, but arbitrary goal-retargeting without further learning is not demonstrated. Neural episodic control provides drastic improvement of learning speed by employing a differentiable memory module with deep learning (Pritzel et al., 2017). (Li et al., 2023) uses instead deep RL with state abstraction. The powerful approach of (Hu et al., 2021) uses DNNs, does not support changes of goals or context, provides no explanations, and does not support implementation with binary synaptic weights. Sequential episodic control (Freire et al., 2025) can provide explanations, but does not demonstrate arbitrary goal retargeting without relearning, and can not be implemented with binary local learning.

Hindsight experience replay enables reuse across goals by relabeling experience for policy or value-function updates (Andrychowicz et al., 2017). EMs are cut there into single transitions, which facilitates reuse but encumbers fast adaptation to new contexts and use of EMs as explanations for action selection. An interesting recent brain-inspired approach to RL, with focus on clever exploration and goal selection, is (Forestier et al., 2022). It may also achieve some form of context-awareness and explainability. But its methods are very different from ours, and it learns continuous vectors of control parameters instead of binary weights. An amalgamation of some of their ideas with ours is likely to enhance both approaches.

There are in addition other methods that enable adaptation to new contexts without relearning. Apparently this had first been demonstrated in (Kaelbling, 1993). More recent methods apply policy-reuse (Rosman et al., 2016) as well as successor features and default representations (Barreto et al., 2017; Piray and Daw, 2021). The meta- and in-context learning approaches of (Zintgraf et al., 2019; Laskin et al., 2022) require DNNs or LLMs. These approaches do not provide explanations through identifiable past episodes.

In most previous methods for using EMs in RL one first learns a value- or Q-function from EMs and then applies greedy action selection with regard to this function. Our method is conceptually different: It demonstrates that it is not necessary to first learn a value- or Q-function. Instead, it uses a brain-inspired shortcut whereby EMs become directly linked to actions via Hebbian learning. These direct links enable action selection with low-latency, since in contrast to value-based approaches the agent does not have to recall the values of all the next states that can be reached by currently applicable actions and compare these values. However, in order to explain the high performance of this simpler approach via direct links, one can show that the stored EMs implicitly define a universal value function, so that action selection via these direct links performs greedy value ascent with regard to this implicit value function. Furthermore, we show that this direct linking of EMs to actions can be learnt through local online learning in shallow neural networks with binary synaptic weights. A RL approach that only requires low resolution of synaptic weights is desirable for efficient implementation in edge devices, e.g. via in-memory computing with highly-energy efficient analog memristors (Li et al., 2017). The brain suggests that RL with low resolution of learnt parameters is in principle possible, since experimental data show that a biological synapse can store at most 5 bits (Bartol Jr et al., 2015; Samavat et al., 2024). In contrast, standard RL methods require a fairly high resolution of parameters that store learnt knowledge.

We show that more detailed information about neural codes that are employed by the brain provides the key for designing algorithms that achieve the previously described combination of functional benefits for RL. Our first observation is that brains employ a contextual encoding of states and objects: A state or object is (also) encoded by the EMs in which they occurred. For example, when we think of our favorite restaurant, we find it hard to do this without automatically recalling also numerous episodic memories of visits to this restaurant. This contextual encoding of states is reminiscent of distributional codes for word meaning had apparently been first proposed by Ludwig Wittgenstein (Wittgenstein, 2009), and which are commonly discussed in linguistics and cognitive science (Lenci, 2018). It is also reminiscent of codes in vector databases in computer science (Taipalus, 2024). Correlations between these vectors provide a measure for the semantic association between two items, e.g, two words. In our more general setting the correlation between the contextual codes for two states, in terms of EMs in which they occurred, provides salient information about their relation. In fact, for navigation in undirected graphs it provides a new type of cognitive map. We show that in contrast to previously considered types of cognitive maps (Behrens et al., 2018; Stöckl et al., 2024; Lin et al., 2026), a cognitive map that results from EMs can be adjusted on the fly to new constraints or contingencies.

The second observation that we take into account is that the brain employs index cells for EMs. For example, there are neurons in the human brain that fire for an extended period of time while the subject watched a specific movie clip, and also during subsequent recall of the same movie clip (Gelbard-Sagiv et al., 2008). The existence of such index cells had already been earlier proposed by (Teyler and DiScenna, 1986). More recent experimental studies demonstrate that such index cells for EMs emerge quite fast in the human brain, after a few repeated presentations of a sequence of images (Kolibius et al., 2023; Purandare and Mehta, 2023; Chettih et al., 2024; Tacikowski et al., 2024; John et al., 2025). In particular, it was shown in (John et al., 2025) that these index cells fire subsequently already for a presentation of a single image of the sequence. Hence, one can approximate the functional role of an index cell for a EM by a Boolean OR over all components this EM (John et al., 2025). On an abstract level one should think of these index cells as pointers to the content of an EM, in particular to neural codes for sequences that can be replayed. These index cells are computationally very useful for retrieving EMs both by a state which occurred in them, and/or by other contextual annotation to EMs as cues.

A third important insight from brain data /is that representations of EMs are strongly coupled with motor areas (Kim et al., 2023). In fact, recall of an EM tends to reactivate motor activities that occurred during the corresponding experience (Boscheron et al., 2026), see also (Robinson et al., 2020). The circuits that implement this coupling are rather complex, involving in particular cortico-basal ganglia-thalamic loops that apparently enable the brain to make the strength of this coupling dependent on reward prediction signals but also on value-free action prediction errors (APEs) (Klaus et al., 2019; Roth and Ding, 2024; Ramot et al., 2025; Greenstreet et al., 2025).

We show that these three observations from the brain provide the basis for an algorithm for goal-directed action selection that strongly differs from previously considered RL methods, but supports the desired combination of brain-like functional benefits: The Episodic Cognitive Map (ECM) algorithm.

The initial pool of EMs consists in the most basic version of the ECM algorithm of random walks. But one can easily boost its performance and simultaneously limit the memory space that it requires through recursive self-improvement. Here older EMs are forgotten and replaced by EMs of subsequent planning solutions, using also the idea of hindsight replay (Andrychowicz et al., 2017).

We explain the basic principles of the ECM algorithm in a generic setting for goal-directed action selection and problem solving: finding a short path from some start node to some goal node in an abstract graph. The ECM algorithm does not require any prior knowledge about the graph. Instead it explores the graph in a self-supervised manner through random walks, and records these experiences in EMs. We show that this yields already close to optimal performance on fairly large random graphs. We then show how one can boost its performance through recursive self-improvement. Finally, we show that the ECM algorithm can also be applied if action outcomes are uncertain, i.e. to Markov Decision Processes (MDPs). We show that it provides also there a more flexible and transparent method for goal-directed action selection, enabling in particular fast online adaption to new goals or contingencies, e.g. safety considerations, and relating chosen actions to concrete experiences that support it.

## 2 Results

### 2.1 Contextual encoding of states

A fundamental concept for goal-directed action selection is the notion of a state. In abstract models for problem solving and navigation in concept spaces, a state is a node in a graph which models this problem solving domain (Sutton and Barto, 2018; Russell and Norvig, 2020), see Fig. 1A for an example. Arguably, the brain employs a contextual encoding of states via episodes in which they occurred. In other words, each state is represented by the web of relations between it and other states that have been created through EMs in which they occurred. On the neural implementation level, this is facilitated in the brain by index cells *z*_*e*_ for episodes *e*, which fire only for those states that were visited during this episode (Teyler and DiScenna, 1986; Gelbard-Sagiv et al., 2008; Kolibius et al., 2023; Purandare and Mehta, 2023; Chettih et al., 2024; Tacikowski et al., 2024; John et al., 2025). An illustration is given for the three episodes depicted in Fig. 1B in Fig. 1C. We model this by an embedding *E* of states *s* into a high-dimensional space that encodes by a binary vector in which Ems this state has occurred, see Fig. 1D. The embedding *E*(*s*) of a state *s* can be trivially computed by a parallel application of all index cells to the state *s*. We refer to Section 4.1 for details of this embedding.

**Fig. 1:**
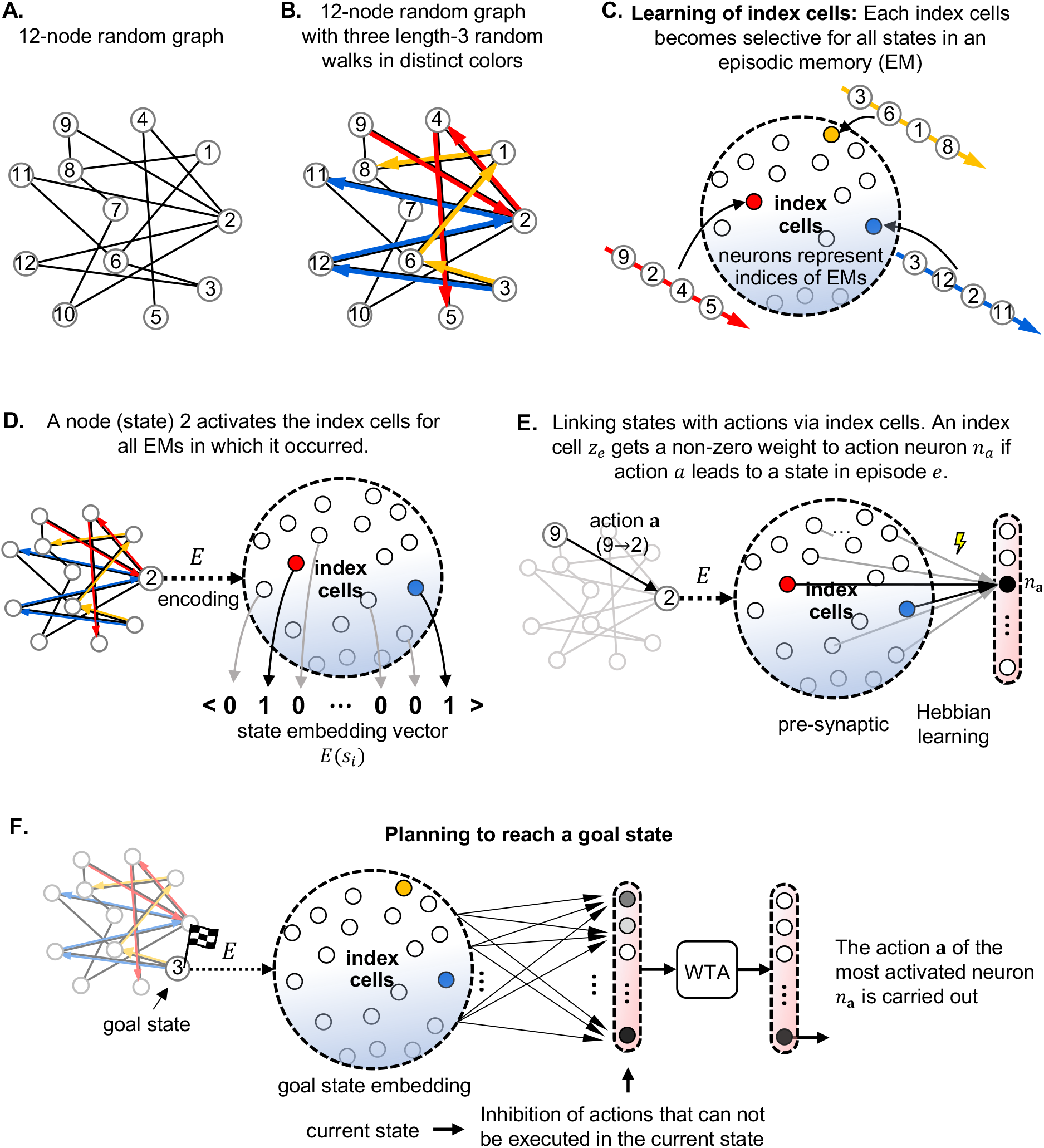
Network-architecture and learning steps for the ECM algorithm. **(A)** A small random graph, which serves as running example for this figure. **(B)** Three sample exploration trajectories indicated by three colors. **(C)** Index cells for these three trajectories (EMs), where each index cell becomes selective for the codes of the graph nodes visited during one of these trajectories. Definition of the embedding *E*(*s*) for the sample node *s* = 2 in the graph. *E*(*s*) is a binary vector which assumes value 1 for each episode in which this node 2 was visited. **(E)** Relating states to actions via the state embedding: Each neuron *n*_*a*_ that can trigger a specific action *a* learns to respond to index cells in the embedding *E*(*s*) of the state *s* that results from executing this action *a*. **(F)** The ECM algorithm for online navigation from the current node to an arbitrarily given goal node *g* = 3 (marked by a flag). One chooses that action which is most strongly activated by the index cells in the embedding *E*(*g*) of the goal state. This is according to the Hebbian learning rule of panel E that action for which the resulting next state co-occurs with the goal state *g* in the largest number of EMs. The next step towards the goal is chosen by iterating this process at the next state.

The index cells for EMs can be created by a simple model for BTSP (Wu and Maass, 2025). It suffices to dedicate just a single neuron *z*_*e*_ to each episode *e*, see Fig. 1C. Note that one cannot replace BTSP in our model by a standard associative learning rule, since this would not capture the random hashing features of BTSP that are functionally important, see the analyses of (Wu and Maass, 2025).

One can make action selection very easy through this contextual encoding of states if the activation of an index cell *z*_*e*_ activates those neurons *n*_*a*_ that can trigger the execution of actions *a* that occurred in the corresponding episode *e*. We model these neurons *n*_*a*_ as linear neurons *n*_*a*_(*x*) for total synaptic input *x*. Activation of these neurons *n*_*a*_ by index cells *z*_*e*_ can be learnt during exploration through one-shot learning with a local rule for Hebbian plasticity with binary weights. This rule sets a weight from 0 to 1 if the presynaptic neuron represents an EM that occurred right after the execution of action *a*; see Section 4.1 for details. It is illustrated in Fig. 1E for the case that its action triggers a move from node 9 to node 2, thereby activating the neurons that fire according to Fig. 1D for the embedding of node 2 into the high-D embedding space. As a result, the neuron *n*_*a*_ becomes activated by a state of the outcome of its action *a*, like mirror neurons in the brain (Bonini et al., 2022; Mukamel et al., 2010). Alternatively, using a slightly different model for EMs which includes the actions that were carried out, one can set the weight from an index neuron to an action neuron to 1 if this action occurred in the corresponding EM; see the last part of Section 4.1. We will show in another paper that this learning process can be understood from a theoretical perspective as application of goal-invariant policy gradient RL (Sutton and Barto, 2018) to an extremely simple neural network. Surprisingly, this simple online learning method is sufficient for producing a short path from any given start node to any given goal node in the task environment, see Fig. 1F. An arbitrary target state *g* (node 3 in this example) is mapped by *E* to the binary vector *E*(*g*), which indicates in which EMs this target state *g* had occurred. We write *A*(*s*) for the set of actions that can be carried out, i.e., is affordable, in state *s*. As a remark on the side we want to mention that in the brain the indirect pathway through the basal ganglia appears to inhibit those actions that are currently not affordable (Klaus et al., 2019). For each candidate action *a*^*′*^, let 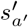 denote the state to which this action leads.

The next action *a* is then selected according to

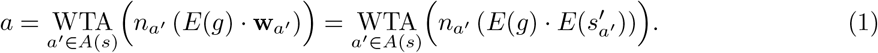

In terms of query-key-value structures, *E*(*g*) serves as the query, 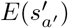 as the key associated with action *a*^*′*^, and the resulting activation of 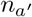 as the corresponding value. We refer to this action selection policy as ECM (Episodic Cognitive Map algorithm) for reasons that are explained in the next Sec. Its strategy is intuitively very clear: It always selects that action for which the state to which it leads occurred in the largest number of EMs in which also the goal node *g* occurred. In case that no such EM exists, heuristic steps such as choosing a random action, or selecting a random state that co-occurred with the goal in an EM as sub-goal, are useful.

### 2.2 The contextual embeddings of states define a cognitive map that guides efficient navigation to distal goals by the ECM algorithm

The quality of an action selection policy is commonly evaluated by the length of paths that it produces for any pair of start and goal states. We consider as example a random graph with 100 nodes, see Fig. 2A, B. We show in Fig. 2C that the ECM algorithm produces paths whose length is close to optimal, provided that it could carry out sufficiently many exploration episodes. This holds in spite of the fact that the ECM algorithm is a low-latency online planning method. Optimal path length can be computed by the offline Dijkstra algorithm.

**Fig. 2:**
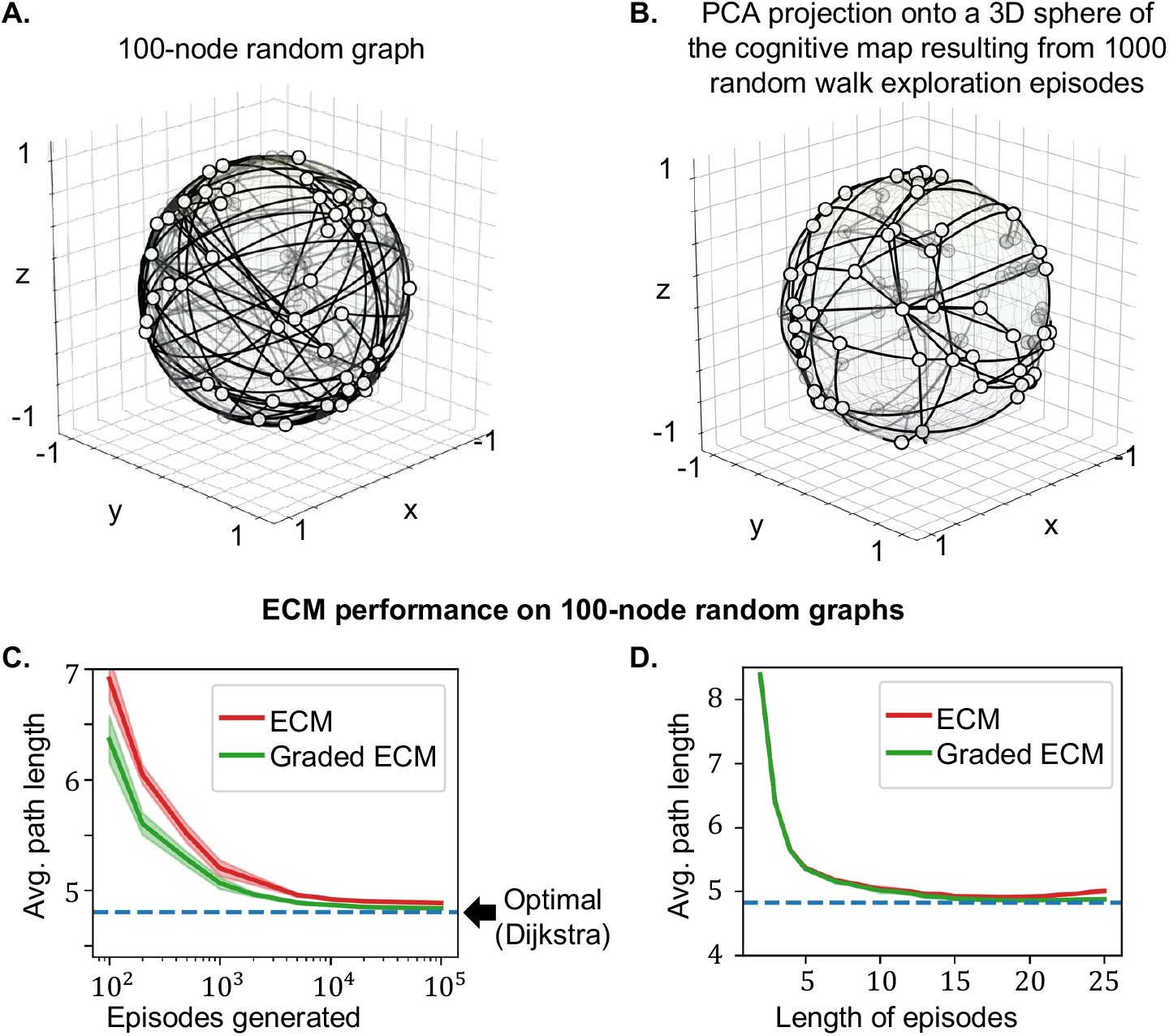
Performance and analysis of the ECM algorithm for action selection in larger general graphs. **(A)** A random graph with 100 nodes and edge probability 3%. Its nodes are plotted at random locations on a 3D sphere. **(B)** Projection of the cognitive map that was created through the contextual embedding of the states according to Fig. 1D onto the same 3D sphere. Distances between embedded states result from correlations between their high-D embedding vectors. They reflect the length of the shortest path between any two nodes. One also sees that moving into the direction of a distal node tends to be a good first step in this cognitive map. **(C)** Average path length for randomly chosen start and goal nodes produced by the ECM algorithm for this graph (red curve). Besides its basic version, we also depict the performance of a variant of the ECM algorithm (green curve) where neurons that provide indices for EMs respond stronger to items close to the middle of the episode (“Graded ECM”). **(D)** Dependence of the average length of produced paths on the length of random walks (EMs) during exploration.

This excellent performance of this simple heuristic raises the question why it works so well. How does it attain the”foresight” that is needed to choose actions that approach the goal even in a non-planar graph? Fig. 2B shows that this is made possible by the way in which relations between nodes are encoded in the geometry of their embeddings (see Fig. 1D) that results from a given set of exploration episodes (1000 random-walk episodes, each of length 16, in this case). As in other types of cognitive maps (Behrens et al., 2018; Stöckl et al., 2024) the geometry of this representation supports a simple heuristic for producing a path to a distal goal that we commonly apply for navigation in 2D environments without obstacles: Choose an action, i.e., an edge in the graph, that moves most directly into the direction of the goal.

Our contextual embedding *E* enables us to apply this simple heuristic also for navigation in random graphs, in spite of the fact that they are in general not planar. As explained in the preceding section, the ECM chooses that action that leads to a state which co-occurred with the goal in the largest number of EMs, i.e., whose contextual encoding is maximally correlated with that of the goal. This strategy can be explained geometrically as moving optimally into the direction of the goal on a hypersphere where contextual codes, encoded by binary vectors (see Fig. 1D), are represented by points on this hypersphere. Correlations between these binary vectors are up to normalization equivalent to the cosine between these vectors. If all nodes of the graph occur approximately in the same number of EMs, this normalization just contributes a common factor to these cosine estimates and can therefore be ignored. Note also that adjacent nodes in the random graph are likely to co-occur in the largest number of random trajectories, and therefore have the smallest distance on the hypersphere. Thus the strategy of the ECM algorithm, choosing a greedy action that maximally increases correlation of the next state with the goal state, amounts to moving on this hyper-sphere into the direction of the goal. Fig. 2B shows that for a graph with 100 nodes the geometry of this cognitive map on a hypersphere is already visible in a projection onto a 3D sphere.

One can also explain the ECM algorithm in terms of a key concept of RL: value functions. The cognitive map that is generated by EMs defines a universal value function *U* (*s, g*) by the correlation between the high-D vectors *E*(*s*) and *E*(*g*) for any states *s* and *g*. The ECM algorithm can then be explained as greedy action selection with regard to this universal value function. But importantly, the ECM algorithm does not have to compute this universal value function for all possible next states, and then select the one of maximal value. Rather, the universal value function is implicitly encoded by the learnt binary synaptic weights from index cells to action neurons, and action selection requires very little computational effort and can therefore be carried out with low latency: For any current state the total synaptic input to each action neuron *n*_*a*_ is equal to the number of episodes *e* that contain both the goal state *g* and the state *s*^*′*^ that results from carrying out action *a* in the current state. Hence, action selection according to Eq. (1) automatically picks an action that leads to a next state of maximal value.

Note that in contrast to most work on learning universal value functions, see e.g. (Schaul et al., 2015; Andrychowicz et al., 2017), our method does not require training of a DNN. SR methods in RL also provide a universal value function without deep learning. But in contrast to SR methods the ECM algorithm only requires binary weights and requires substantially less computational effort for action selection. We will see in the next sections that additional differences between the ECM and SR algorithms are that the ECM algorithm provides explanations for its action selections, and its action selection can be adjusted on the fly to new contingencies. Furthermore we will show in Fig. 6 that learning requires on a digital computer for the ECM algorithm substantially less computational effort than for the SR algorithm.

Fig. 2C shows that the performance of the ECM algorithm can be somewhat improved by a refined action selection strategy to which we refer as graded ECM, that aims at reaching next states that are connected by short paths with the goal. The only new feature of the graded ECM algorithm is that it employs index cells that respond in a bell-shaped manner to the nodes in the episode which they encode, most strongly to nodes near the center of the episode. Such a response profile of index cells is actually suggested both by experimental data on BTSP, see e.g. Fig. 3E of (Milstein et al., 2021), and by recordings from the human brain, see Fig. 2 of (John et al., 2025). A simple way to mimic this biological detail in a model is to let the neuron that encodes a sequence according to Fig. 1C respond stronger to items near the middle of the sequence. This implies that the nodes of the graph are mapped by the embedding *E* to analog rather than binary vectors. Then the action selection mechanism of Fig. 1F prefers actions where the next state is connected by shorter episodes with the goal node. These shorter episodes have a larger chance that either the next state or the goal state is closer to the middle of the episode, thereby contributing more to the correlation between the corresponding vectors of analog values. The green curves in Fig. 2C, D show that the resulting “graded ECM” algorithm improves performance in the case of relatively few exploration episodes, see Methods for details.

**Fig. 3:**
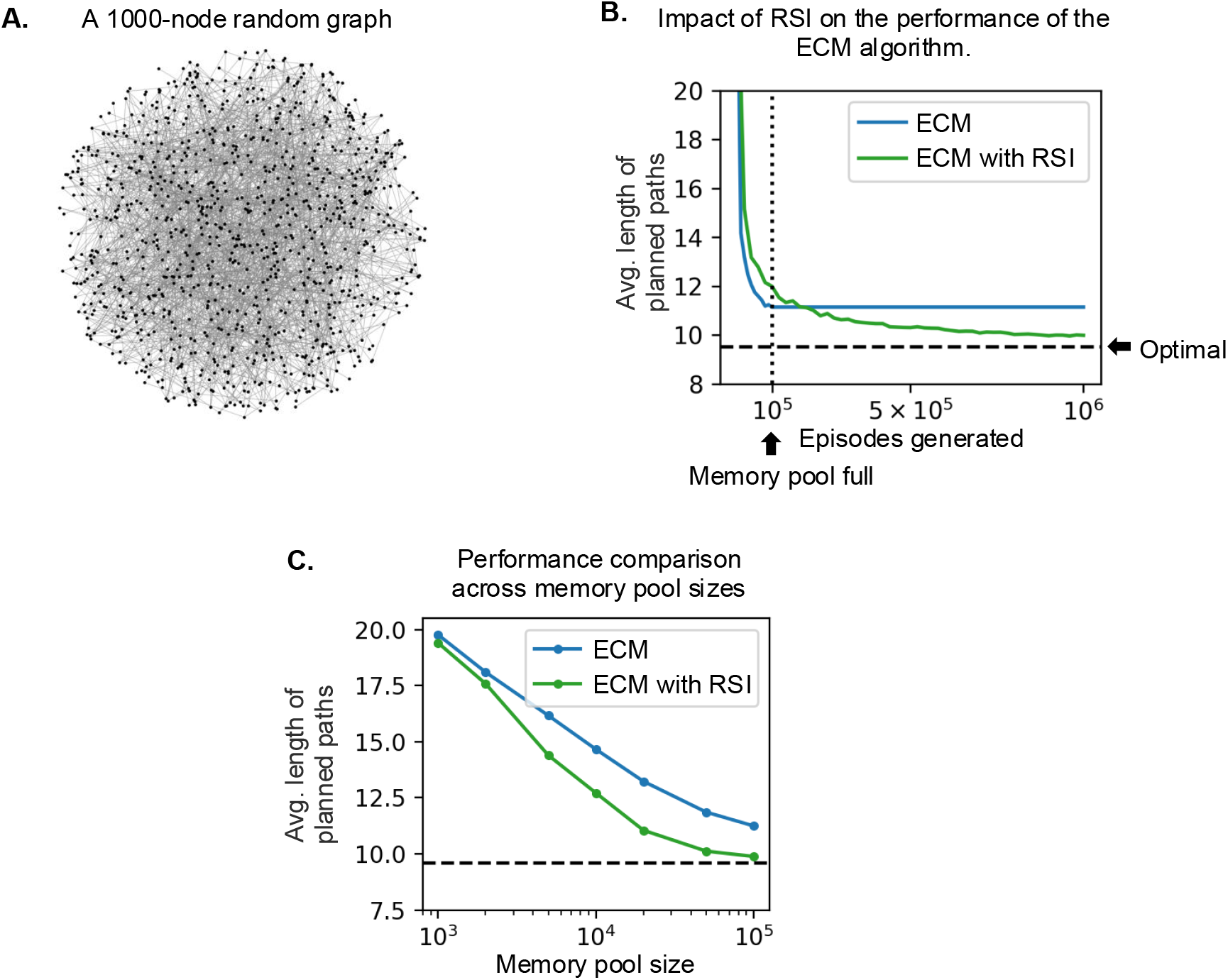
Impact of RSI on the ECM algorithm for navigation in a 1000-node random graph. **(A)** The 1000-node random graph used in the experiment. Node locations are uniformly sampled within a circle. **(B)** Average planned path length as more episodes are generated during exploration. The dotted vertical line marks the point where the size of the memory pool has reached a given ceiling. The ECM algorithm with RSI keeps improving after this point, whereas the basic ECM cannot do so. **(C)** Performance comparison across memory pool sizes. The ECM algorithm with RSI can produce shorter planned paths than the basic ECM algorithm for many different sizes of its pool of EMs.

### 2.3 Forgetting of EMs and recursive self-improvement of the ECM algorithm

The brain employs sophisticated mechanisms for evaluating EMs and forgetting less useful ones. Analogous methods have also been developed in machine learning, see e.g. (Steunebrink et al., 2016). Both sources provide numerous ideas for improving the ECM algorithm. We present here one such advanced version that we call the ECM algorithm with RSI because it adds Recursive Self-Improvement. The main idea is to create index cells *z*_*e*_ not only for trajectories that arise during initial exploration, but also for some paths *e* which the ECM algorithm produces in subsequent task applications. Naive first-in, first-out (FIFO) replacement, in which each newly generated episode replaces the oldest EM regardless of its usefulness, does not show the desired recursive self-improvement. We suspect this is because when a suboptimal episode is admitted to the memory pool, it can further bias subsequent action selection toward inefficient routes. Storing further episodes generated by these choices can then reinforce the same behavior, even after the original episode has been forgotten. We therefore evaluate the usefulness of candidate episodes and retain them selectively.

To decide whether an index cell should be created for such a path, the ECM algorithm with RSI stores for each pair *a, b* of nodes the length of the shortest path from *a* to *b* that it has found so far. It then examines for the new path *e* whether a sub-path of it provides for some nodes *a*^*′*^ and *b*^*′*^ a shorter path from *a*^*′*^ to *b*^*′*^ than found so far. If that is the case, it creates an index cell for *e*. Simultaneously it forgets that EM, or more precisely deletes the index cell for that EM that currently appears to be least useful in terms of the lengths of path segments that it provides, see Methods. Deleting an index cell also removes its outgoing connections to action neurons. For the new index cell, these binary connection weights are set to 1 for actions leading to states contained in the new episode, and to 0 otherwise. It does this also if the new path *e* does not reach the goal that it was meant to reach, thereby implementing an idea of the hindsight replay method from (Andrychowicz et al., 2017). However, no replay of stored episodes is needed by the ECM algorithm.

The resulting recursive self-improvement character of the ECM algorithm is clearly visible in Fig. 3B: Its performance keeps improving even after a fixed size memory pool is full. Fig. 3C shows that the ECM algorithm with RSI performs better than the basic ECM algorithm for almost any pool size.

Note that one can trivially expand the previously described method to the case where no EM contains a path from the current start *a* to the given goal *b* by considering other states in paths that visited *b* as possible sub-goals, or by carrying out further exploration. Furthermore, for applications in environments that have in contrast to random graphs an inherent structure, one can use inspiration from the brain for evaluating pairs of episodes as being more or less similar, and one can consolidate similar episodes into a more general scheme, see e.g. (Stickgold and Walker, 2013). These examples show that there are many ways for improving the ECM algorithm, some of them inspired by corresponding processes in the brain.

### 2.4 The ECM algorithm enables context-aware decision making

Besides its inherent flexibility to changes of the goal and its arguably minimal computational complexity, the ECM algorithm provides two further functional benefits: Context-awareness of action selection and explainability of action selection (the latter will be addressed in Section 2.5). These benefits arise from the fact that the ECM algorithm does not compress all learnt knowledge into parameter values, but retains -like the brain-salient original experiences (EMs). The brain evaluates and annotates EMs online from numerous behaviorally relevant perspectives. According to (Eichenbaum, 2017) this is carried out by several brain areas that are interconnected with area CA1. These internal evaluations of EMs are used by the brain to selectively recall only those EMs that have a number of particular qualities. For example, we can recall selectively only EMs of particularly pleasant visits to a certain restaurant, or only those on rainy days, only those where we were stuck with the bill, or only those EMs that satisfy a combination of such properties.

One can easily port this enormous versatility of memory recall by our brain into models by adding a context- and evaluation module to our model, shown symbolically at the top of Fig. 4A. This module enables the ECM algorithm to selectively activate only those EMs that contain the goal, and which have in addition other properties that arise from the current task context. Since the episodic cognitive map, and its associated universal value function, arise from the currently activated set of EMs, this allows an online adjustment of both to the current task context. In order to avoid confusion, we would like to add that this resulting context-awareness of decision making is unrelated to the contextual encoding of states, which just referred in general to the temporal context of states in EMs .

**Fig. 4:**
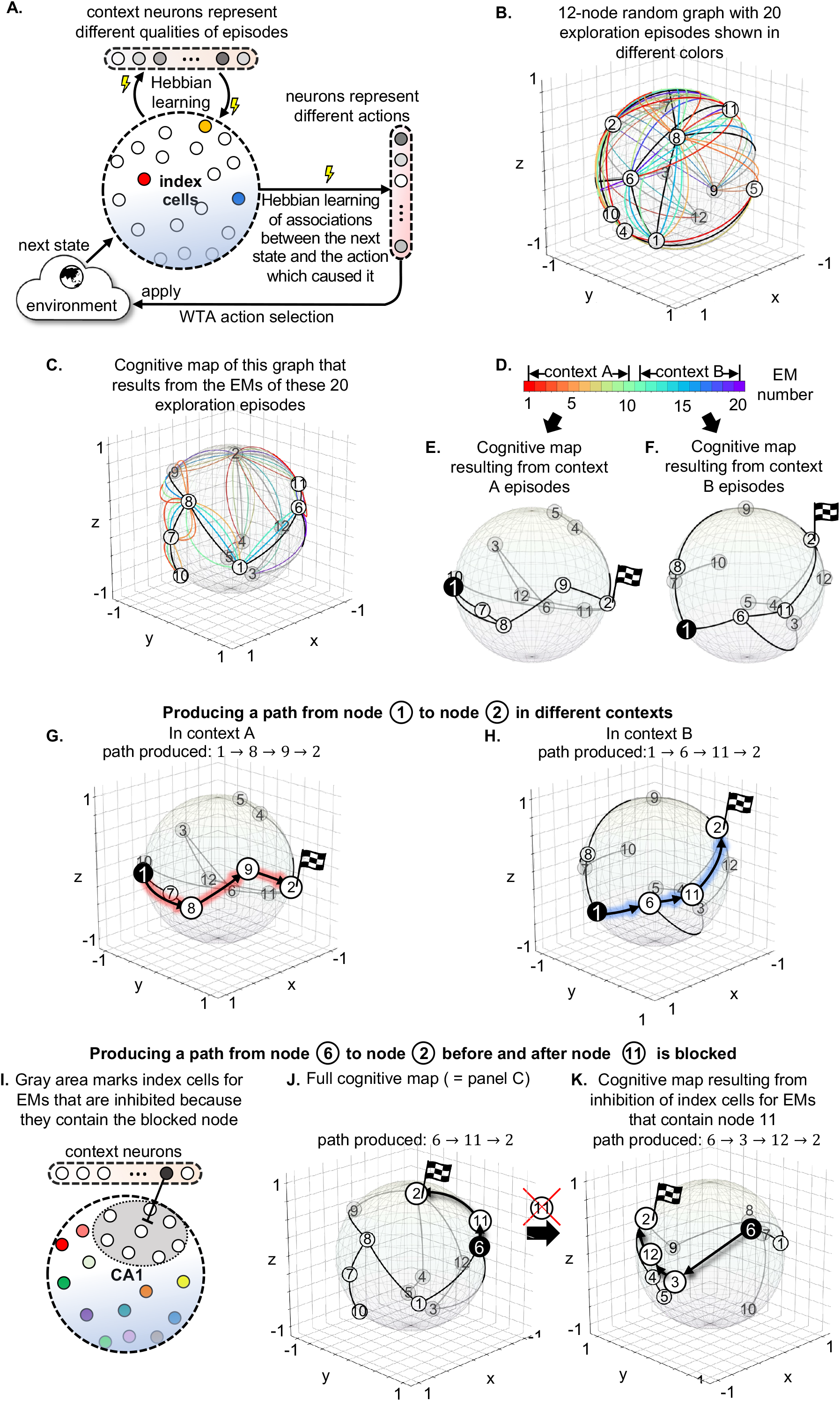
Context-aware action selection through the ECM algorithm. **(A)** Augmentation of the architecture through a network module (shown at the top) which evaluates the quality of EMs from various perspectives. Synaptic connections from this evaluation module to CA1 can selectively activate or inhibit EMs according to the current context. **(B)** The graph from Fig. 1A, with 20 exploration episodes shown in different colors. **(C)** The cognitive map for this graph that results from the embedding according to Fig. 1D into index cells for these 20 EMs. For simplicity we are labeling here and in the following panels the embedded nodes of the graph by the same label as the corresponding nodes. **(D)** An arbitrary partition of the 20 episodes into two contexts A and B. **(F)**Projection of the cognitive map that results from the 10 episodes in context A onto a 3D sphere. Same for context B. **(G, H)** The task to produce a path from node 1 to node 2 yields two different solutions in contexts A and B, indicated via projections of the associated cognitive maps onto a 3D sphere. **(I)** Another context (constraint): Node 11 can currently not be used for navigation in the graph. The cognitive map is automatically adjusted to this new constraint by inhibiting index cells for EMs that contain node 11. **(J, K)** Task: Generate a path from node 6 to node 2. Panel J shows the cognitive map before removal of node 11, and panel K shows the instantaneous update of the cognitive map after removal of node 11. Note that the removed node 11 lies on the shortest path for this task.

We demonstrate the inherent capability of the ECM algorithm to make action selection context-aware in Fig. 4 for navigation in the simple graph from Fig. 1. 20 exploration episodes are marked with different colors in panel B, and partitioned arbitrarily into two contexts A and B according to their color, see panel D. Note that the embeddings *E*(*s*) of states (nodes) *s* are labeled with the same number as the state (node) *s*. Each of the two contexts yields a different cognitive map. These are visualized in panels E and F via projections onto a 3D sphere, like in Fig. 2B. One sees that the task to produce a path from node 1 to node 2 yields in these two different contexts completely different solutions, see panels G and H. The beauty is that this specialization of the cognitive maps to a given context does not require any computational overhead or new learning: One just has to constrain the activation of EMs by the current goal node to those EMs that fit into the desired context, as signaled by the evaluation module. Also, if the task environment suddenly changes and some nodes are no longer available as transition points, e.g. if a road is blocked in a spatial navigation task, the ECM algorithm can inhibit on the fly those EMs which contain these nodes, as demonstrated in panels I, J, K of Fig. 4.

One can also take into account how recently an EM was acquired, and use that information to solve RL tasks with drifting reward probabilities, e.g. for multi-armed bandits (Bornstein et al., 2017). For that one needs a memory control module that activates index cells for more recently rewarded EMs more strongly than others that contain the same goal of getting a reward. These index cells have then automatically a stronger impact on action selection according to the selection strategy of the ECM algorithm, see Fig. 1F.

There exist numerous methods for making RL context-aware, see e.g. (Finn et al., 2017; Achiam et al., 2017; Christiano et al., 2017; Chen et al., 2021). But most of these methods require offline training through stochastic gradient descent in DNNs or LLMs. In contrast, the context-aware decision making of the ECM algorithm arises from local online learning rules, and can be implemented by a shallow neural network.

### 2.5 The ECM algorithm provides efficient, context-aware, and explainable action selection also when action outcomes are uncertain

The same ECM algorithm can also be applied if it is uncertain to which next state (node) an action leads, i.e., for Markov Decision Processes (MDPs), see Fig. 5A. To be precise, the standard definition of a MDP includes the specification of a reward function. But since the learned representation of the ECM algorithm does not depend on the specification of a particular goal, MDPs without a fixed reward function are a more suitable application domain. We denote these as MDP^-R^s. In (Liu et al., 2024) the term controlled MDP was used for them, but this term might be confusing in our context.

**Fig. 5:**
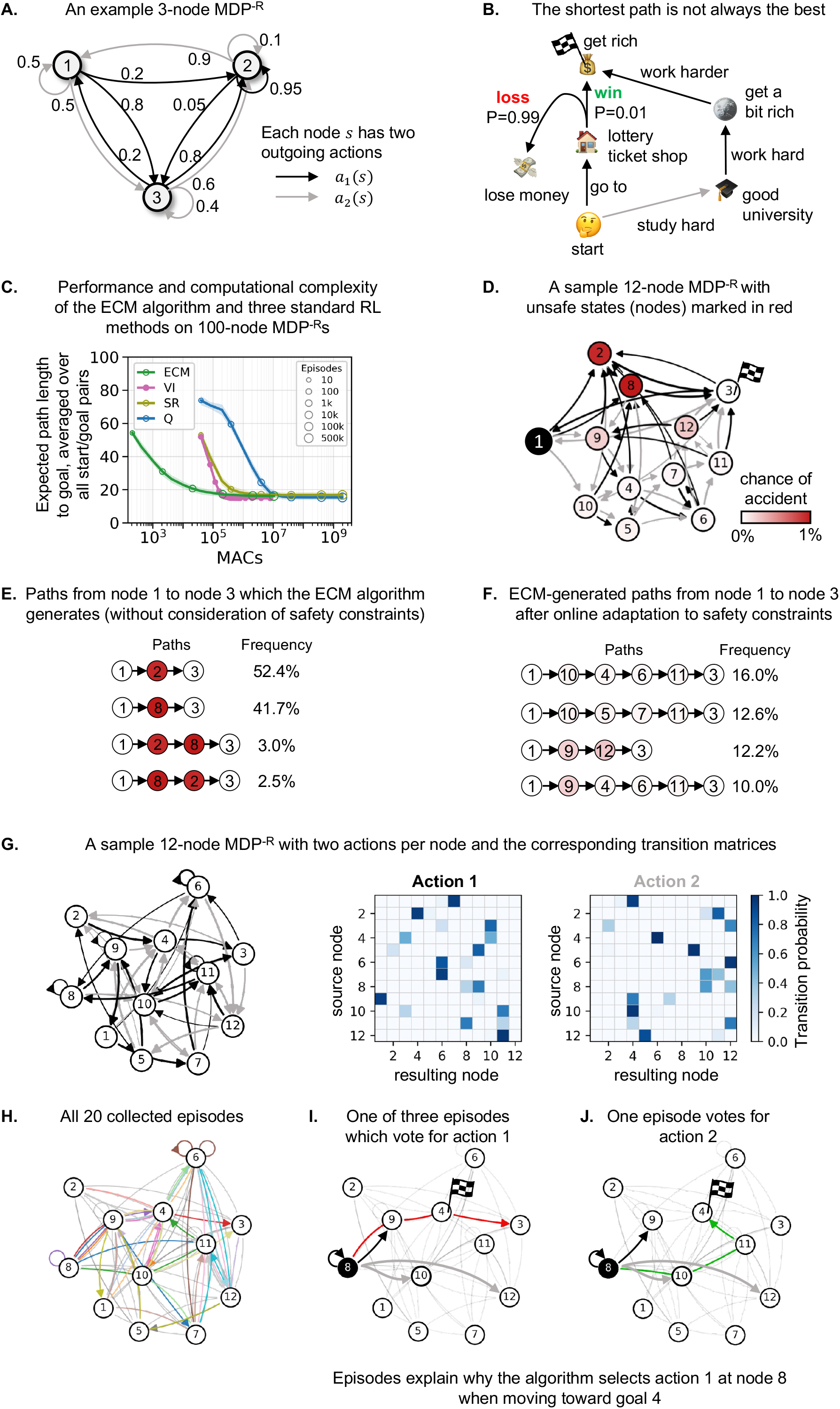
Efficient context-aware and explainable action selection of the ECM algorithm for navigation in MDP^-R^s. **(A)** A sample 3-node MDP^-R^ with *A* = 2 actions (black and gray) at each node, each leading with certain probabilities to *K* = 2 different next states. *A* and *K* are fixed across the examples in this figure, although the ECM algorithm can be applied for any values of them. **(B)** The “get rich” example demonstrates the trade-off between short but uncertain action sequences and longer but more reliable ones. **(C)** Performance of the ECM algorithm, VI, SR, and Q-learning in terms of expected path length, averaged over all ordered start–goal pairs and 10 independently generated 100-node MDP^-R^s instances, as a function of the computational cost invested into learning. The latter is measured by the number of multiply-accumulate (MAC) operations applied during learning. Shaded areas indicate *±* one standard deviation across the 10 MDP^-R^s instances. For VI, data points show performance after selected complete value-iteration sweeps; for ECM, SR, and Q-learning, they show performance after selected cumulative numbers of random-walk exploration episodes. The horizontal coordinate gives the cumulative learning cost at each checkpoint. See Sec. 4.7.6 for more details. **(D)** A 12-node MDP^-R^ in which each node is assigned a risk (chance of accident). Node color indicates the risk probability. Node locations are manually placed in this plot. **(E)** Paths from node 1 to node 3 generated by the policy of the ECM algorithm without safety constraints. **(F)** Paths from node 1 to node 3 generated by the ECM algorithm, with each index cell assigned a graded activation according to the overall safety of its corresponding episode. Index cells associated with safer episodes receive higher activation. **(G)** An example 12-node MDP^-R^ defined by the transition matrices on the right. Each node has two outgoing actions, shown in black and gray. Node locations are manually placed. **(H)** 20 exploration episodes (EMs). **(I)** One of the three EMs supporting the choice of action 1 at node 8 when navigating from node 8 to node 4. **(J)** The single EM supporting the non-chosen action 2 at node 8. Note that it contained a longer path from 8 to 4 than the EM in panel I that voted for action 1. Panels I and J explain why the agent selects action 1 at node 8.

Goal-directed action selection in the presence of stochastic action outcomes requires even more sophisticated foresight than in the deterministic case, because a new tradeoff arises between short but uncertain sequences of action selections for reaching a given goal and longer but more reliable sequences. Fig. 5B illustrates this tradeoff for a hypothetical task where the goal is to get rich. In this case the longer but safer path resulting from studying and working hard has to be compared with the shorter but less certain path via purchase of a lottery ticket.

Note that for navigation in MDP^-R^s the ECM algorithm has to produce for a given goal not just a short path from the start to the goal, but a policy that provides an action selection for any node in the graph that might become a transition point. The quality of such a policy is commonly measured by the expected length of the resulting path to the goal, averaged over all pairs of start and goal nodes.

One other inherent difficulty of navigation in MDP^-R^s is that state transitions are in general asymmetric, i.e., actions can not necessarily be inverted. Actually, in view of stochastic action outcomes it is not even clear what such an inversion would mean. This asymmetry also affects the use of cognitive maps for navigation: One can no longer base navigation on reduction of the distance to the goal, since a distance metric is inherently symmetric.

In spite of these new difficulties, one can basically use the same ECM approach as for deterministic symmetric graphs. The only difference is that the synaptic weight from an index cell *z*_*e*_ to an action neuron *n*_*a*_ assumes here the value 1 only if *a* is the first action that is executed in episode *e*. This makes sure that an action selection cannot be erroneously induced by an EM in which the chosen action occurred after the currently given goal was visited. If one has too few EMs one can formally split a single episode *e* it into several EMs, each starting at a different action in *e*.

One can then use exactly the same action selection mechanism as for undirected deterministic graphs. The important difference is that for an MDP^-R^ there is no unique state 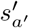 to which an action *a*^*′*^ leads. Instead, the incoming weight vector **w**_*a*_*′* of action neuron *n*_*a*_*′* indicates which exploration episodes were initiated by action *a*^*′*^. The next action is therefore selected according to

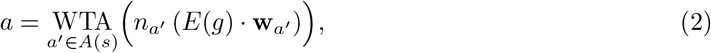

Thus, the ECM algorithm chooses that action for which exploration episodes that started with this action had reached the goal most often. The statistics of these episodes defines a universal value function *U* (*s, g*) for arbitrary pairs of states *s* and goals *g*. For MDP^-R^s it is in general asymmetric, i.e., in general *U* (*s, g*) ≠ *U* (*g, s*). The ECM algorithm approximates greedy action selection with regard to a slight variation of this universal value function (where one normalizes *U* (*s, g*) by the number of visits to node *s* during exploration). A nice feature is that the ECM algorithm never has to compute any value function, which makes action selection fast and efficient; see Sec. 4.6.2 in Methods for details.

A gold standard for action selection in MDP^-R^s is more difficult to compute since in this case the Dijkstra algorithm does not produce policies with expected shortest path lengths. But optimal policies can be computed via iterative RL methods such as value iteration (VI) (Sutton and Barto, 2018). We show in Fig. 5C that the ECM algorithm produces approximately optimal policies. In addition it has the advantage that it produces already reasonable solutions with a smaller number of computation steps during learning. This holds in spite of the fact that the ECM algorithm is only provided with exploration episodes, whereas in our comparison a perfect model of the MDP^-R^ is given to the VI algorithm without any charge. Since VI has to be restarted from the beginning when the goal changes, the computational cost of it is given in Fig. 5C for the total amount of computations during learning when VI is applied sequentially for any possible node as goal. We also compare there the computational cost with that of Q-learning.

The SR variant of RL (Dayan, 1993) also learns a universal value function approximator. However, SR compresses all episodic experiences into expected state-occupancy values, without keeping track of the individual EMs that support it. This difference endows the ECM algorithm with three functional advantages over SR methods: Its universal value function can be adjusted on the fly by inhibiting index cells for EMs that do not fit to current constraints or other context information, those EMs that support a concrete action selection can easily be recovered, and it requires substantially less computational effort for learning, see Fig. 5C.

Fig. 5C shows that ECM learning is from the perspective of computational complexity more efficient than for VI, SR, and Q-learning. We demonstrate the capability of the ECM algorithm to adjust on the fly to new contingencies in Fig. 5D–F for an example. Assume that new information about the risk involved in passing through some states in a MDP^-R^ has suddenly become available. If context neurons (symbolically shown at the top of Fig. 4A) inhibit index cells for EMs *e* more or less, according to the risks of nodes which they visit, then EMs that involve higher risks have according to the scheme of Fig. 1F lower impact on action selection (see Sec. 4.7.3); in other words, episodes that encountered lower risks contribute more strongly to action selection. This shifts in the example of Fig. 5D the resulting policy from the risk-agnostic policy that only aims at short paths, as indicated in Fig. 5E, to the safer policy whose generated paths are shown in Fig. 5F. Importantly, this adjustment requires no relearning of the stored episodes or action associations; only the contribution of the existing EMs to action selection is changed.

As in the deterministic case, each action selection of the ECM algorithm in a MDP^-R^ is supported by a concrete set of experiences (EMs), which provide an explanation for its selection, see Fig. 5G– J for a visualization in the case of a small instance of the problem, shown in Fig. 5G. One has at each node a choice between two actions, indicated by black and gray arrows (each action with two possible outcomes). The outcome probabilities are given by the matrices on the right. For ease of visualization, consider a case where only the 20 EMs depicted in Fig. 5H are provided for action selection. We consider in Fig. 5I the case that the black action has been chosen at node 8 in order to reach node 4. This choice is supported by three prior experiences (exploration trajectories), only one of them is depicted here for simplicity. In order to recover these episodes that explain an action selection one only needs to complement the synaptic connections from index cells to action neurons, see Fig. 1E, by synaptic connections in the inverse direction that carry the same weight. Then the activation of an action neuron *n*_*a*_ activates the index cells for those episodes that support its choice.

Numerous methods for explainable RL have already been proposed, see the review (Milani et al., 2024). Many types of explanations involve features of states that are salient for action selection. These types of explanations are not possible for navigation tasks in MDP^-R^s, since no information about features of nodes is provided there. But in addition to direct explanation for an action selection, also counterfactual explanations can be generated by the same mechanism by briefly activating the action neuron for a non-chosen action, see Fig. 5J. Since both types of explanations emerge automatically from the shallow network that the ECM algorithm employs, the ECM algorithm falls according to the taxonomy of (Milani et al., 2024) into the category of intrinsically interpretable action selection strategies.

Note that in the brain both index cells for EMs and neurons that can trigger the release of an action that occurred in it are likely to be part of a loop that interconnects cortical areas with the basal ganglia and the thalamus (Klaus et al., 2019). These loops could support both triggering of an action on the basis of EMs as in (Boscheron et al., 2026), and the inverse of it, i.e., recover EMs that involve a specific action.

### 2.6 Scaling of performance and computational complexity of the ECM algorithm

We tested the performance and computational cost of the ECM algorithm on substantially larger and more complex MDP^-R^s than those in Fig. 5: MDP^-R^s with 1000 nodes and three actions available at each node (Fig. 6). We compared the ECM algorithm again with the same three common RL approaches as in Fig. 5: VI, SR, and Q-learning.

**Fig. 6:**
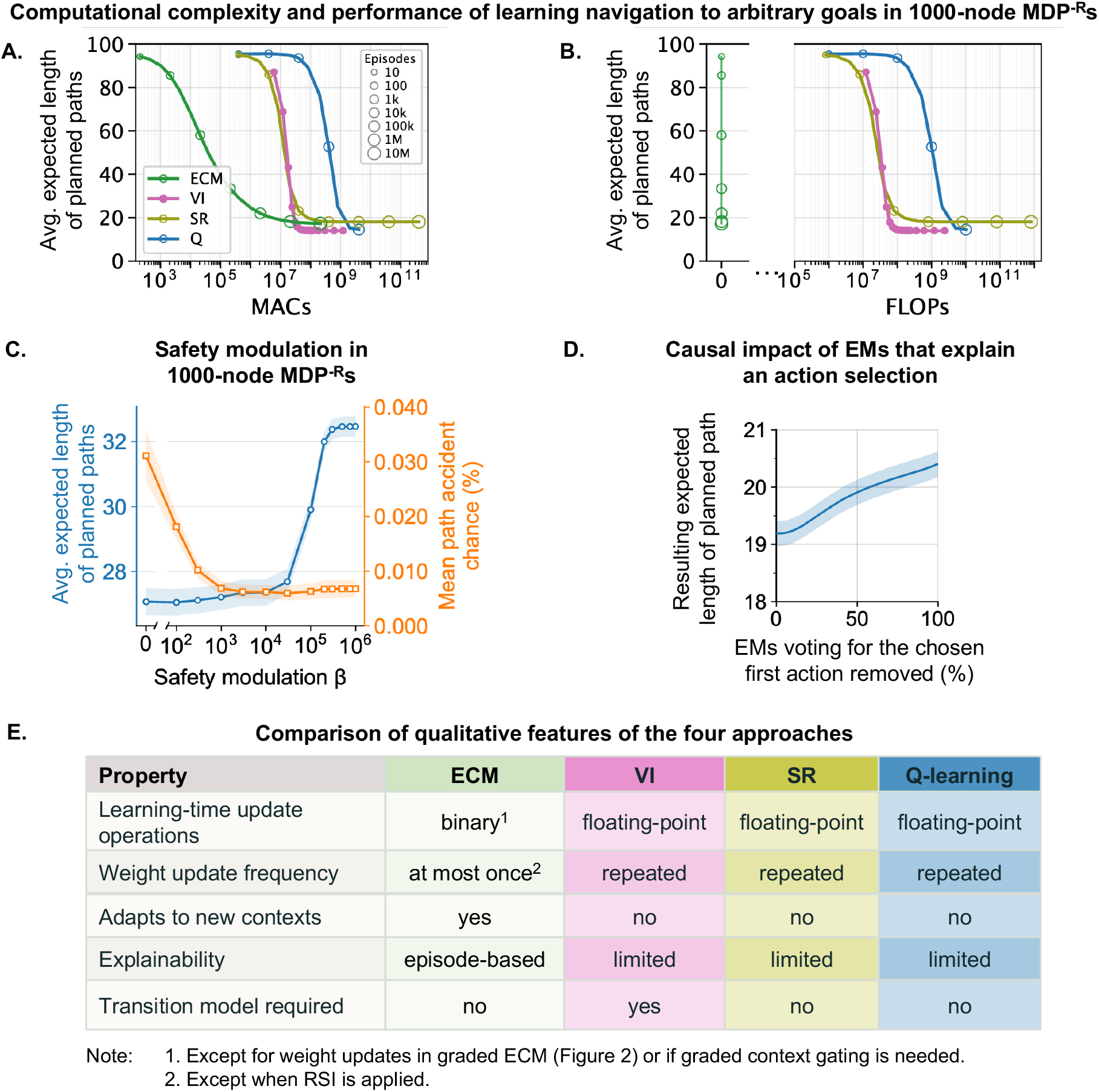
ECM combines efficient learning with flexible and transparent action selection in larger environments (1000-node MDP^*−*R^s with three actions per node). **(A)** Average expected path length for arbitrary start and goal states versus multiply–accumulate operations (MACs) used for learning. ECM achieves short expected path lengths with substantially fewer learning MACs than VI, SR, and Q-learning. Conventions as in Fig. 5C. Panels A,B show the mean *±* standard deviation (shaded) across 10 randomly generated MDP^*−*R^ instances; standard deviations are smaller than in Fig. 5C and not visible here. ECM reaches near-optimal expected path length after approximately 10^6^ exploration episodes of length *L* = 20. Data points show performance after selected numbers of complete value-iteration sweeps (VI) or random-walk exploration episodes (ECM, SR, and Q-learning). In A,B, horizontal axes show the corresponding cumulative learning cost; marker sizes indicate episode counts, except for VI. Each VI data point represents one value-iteration round over all nodes in the value tables for all goals. See Sec. 4.7.6 for details. **(B)** Performance versus floating-point operations (FLOPs). ECM’s learning rule requires no additions or multiplications because episodic memories and action associations are stored through binary memory writes. **(C)** Safety modulation in ECM. Average expected path length (blue, left axis) and mean path accident probability (orange, right axis) versus the safety modulation parameter *β*. Increasing modulation initially reduces accident probability; stronger modulation lengthens paths without further safety gains. **(D)** Effect of removing episodes supporting ECM’s first action for a given start–goal pair. Average expected path length versus the percentage of these voting episodes removed reveals an almost immediate causal effect on the chosen path (policy). **(E)** Table summarizing qualitative differences among the four algorithms. ECM uses binary weights, writes each weight at most once, adapts to new contexts, explains action selection in terms of episodes, and requires no pre-specified state transition model.

Panel A shows that the ECM algorithm reaches short expected path lengths with substantially fewer multiply-accumulate operations (MACs) during learning than VI, SR, and Q-learning. MACs have been proposed as a useful measure of computational complexity for neural-network implementations (Sze et al., 2017). Since a MAC combines memory access with arithmetic operations, we show in panel B a special advantage of ECM that its learning requires no floating-point additions or multiplications (under our operation-counting convention, see Methods). Note that VI has a somewhat unfair advantage in this analysis of learning complexity, since it is given the perfect model of the MDP^-R^ for free, while the other algorithms only access exploration episodes that are provided (without charge for their generation). Since the evaluation has a fixed maximally allowed steps (we set it to 100), trajectories that have not yet reached the goal could lead to an underestimation of the average path length. We therefore show in Fig. S2 of Supplementary Information, Sec. C how often the goal is reached within this horizon. These success rates approach 100% as learning converges, which guarantees the horizon setting is fair in this regime. We also tested whether using just one rollout for each start–goal pair affects the comparison. Fig. S3 shows that the performance curves remain nearly unchanged when 10 rollouts are used for each pair.

We next examined whether context-dependent action selection also scales to these larger environments. We assigned accident probabilities to a sparse subset of nodes and varied the safety modulation parameter *β*, which controls how strongly less safe episodes are suppressed during action selection, while keeping the stored EMs and action associations fixed (Fig. 6C). Increasing *β* from 0 to 10^3^ reduced the mean path accident probability from approximately 0.031% to 0.0069%, making paths about 4.5 times safer, while average path length increased only slightly, from 27.1 to 27.2 transitions. Thus, without relearning, ECM can flexibly trade off safety against a small increase in path length.

The same access to individual EMs also enables ECM to explain its action choices in terms of concrete prior experiences. We evaluated the faithfulness of these episode-based explanations by progressively removing the contributions of the identified EMs that voted for the first action, and recomputing the first action; for a generic but fixed start-goal pair (Fig. 6D). As more supporting EMs were removed, the path length increased, from approximately 19.2 to 20.4 transitions at complete removal, with subsequent decisions using all remaining EMs. Thus, the originally selected first action depended on the identified supporting EMs that can be viewed as explanations for the first action.

Panel E summarizes other major differences between the four algorithms for navigation in MDP^-R^s. The absence of floating-point arithmetic during ECM learning is relevant to hardware architectures that support local synaptic storage and updates, including SRAM-based digital neuromorphic processors (Frenkel et al., 2018), magnetic tunnel junction synapses (Zhou et al., 2025), and memristive arrays (Kim et al., 2021). In the basic ECM algorithm, each binary weight undergoes at most one logical 0 *→* 1 transition before its memory slot is reused. This could enable simple local write control and reduce transfers between memory and a separate arithmetic unit. In memristive implementations, binary weights could also simplify programming compared with precise tuning of multilevel conductances, which can require repeated write–verify operations (Kim et al., 2021; Mackin et al., 2022; Song et al., 2024). These properties motivate potential hardware efficiencies, while actual energy savings depend on memory capacity, physical programming costs, peripheral circuitry, and episode replacement.

## 3 Discussion

For embodied AI, especially for edge devices, we need light-weight methods for goal-directed action selection that are transparent and can quickly adjust to new goals and constraints. Furthermore, edge devices should be able to acquire necessary knowledge about their environment and the consequences of own actions through online learning in shallow networks, ideally through on-chip learning on neuromorphic chips. In addition, low latency of decision making is essential. There exist already many partial algorithmic solutions that satisfy some of these constraints. But the brain shows that in principle all of them can be satisfied by a single system. We have shown that experimental data from the human brain provide useful hints for porting similar capabilities into artificial systems. In particular, contextual encoding of states, index cells for EMs, and direct coupling of EMs to motor-activity provide the basis for a previously not considered strategy for learning goal-directed decision making, the ECM algorithm. Since index cells can be viewed on an abstract level as pointers to the content of EMs, and since the brain-inspired algorithm is able to consult many of them in parallel in order to select an action, it provides an interesting paradigm for efficient parallel computation in pointer architectures.

The ECM algorithm retains and uses individual experiences for action selection. This places the ECM algorithm into the tradition of other exemplar based methods in machine learning, such as k-NN, Support Vector Machines, and case based reasoning. The ECM algorithm also inherits the inherent transparency of these exemplar-based methods: Decisions can be explained by concrete exemplars that support them. In the case of the ECM algorithm, each action selection can be explained in terms of specific EMs that contributed to it, and also counterfactual explanations can be provided, see Fig. 5I,J. Another inherent benefit of the exemplar-based approach of the ECM is that it can adjust its action selection on the fly to a new context by relying only on those EMs that are consistent with this new context, see Fig. 4. Furthermore, the ECM algorithm can autonomously improve its knowledge base from an initial exploration phase through a form of recursive self-improvement where records of solved planning tasks provide additional EMs. We have shown that one can use here a similar trick as in (Andrychowicz et al., 2017) to extract useful information also from unsuccessful planning attempts. In addition, the relative usefulness of individual EMs can be autonomously examined by the algorithm. In this way, the size of the pool of retained EMs can be improved and simultaneously reduced in size, see Fig. 3B, C. These offline examinations of EMs are reminiscent of replay phases in the brain (Chen and Wilson, 2023).

Many RL algorithms, such as temporal difference (TD) learning, have been motivated by earlier experimental data which showed that phasic increases of dopamine signals in the brain are related in some behavioral tasks to reward-prediction errors (Gershman et al., 2024). More recent experimental data have challenged this interpretation, since dopamine was found to serve in other experimental settings as value-free teaching signal, that is for example related to curiosity, surprise, and action prediction errors (Sharpe et al., 2020; Vellani et al., 2020; Jeong et al., 2022; Markowitz et al., 2023; Lee et al., 2024; Costa and Schoenbaum, 2025; Greenstreet et al., 2025). Our results suggest that EMs support efficient goal-directed navigation also in the absence of reward-prediction signals. Furthermore, they are consistent with an alternative interpretation of phasic increases of dopamine that has been proposed in a number of experimental studies: as signal that promotes the formation and retention of episodic memories for salient events (Adcock et al., 2006; Shohamy and Adcock, 2010; Bethus et al., 2010; Poh et al., 2022). Such a saliency-driven filter for the retention of EMs would enable the ECM algorithm to operate with smaller pools of EMs.

We have shown that the high performance of the ECM algorithm in environments where actions are reversible can be explained by a new type of cognitive map that is formed in an online manner through relations between contextual encoding of states, see Figs. 2 and 4. A major functional advantage over the type of cognitive maps considered in (Behrens et al., 2018; Stöckl et al., 2024; Lin et al., 2026) is that these cognitive maps do not have to be learnt and represented explicitly. This has, apart from the ensuing enhancement of the efficiency of learning and computing, the advantage that this type of implicit cognitive map can be reconfigured on the fly if constraints or contingencies change that cause a focus on a particular subset of EMs, see Fig. 4D–K and Fig. 6C. Furthermore, we have shown that for navigation in directed graphs and MDPs, where a symmetric distance metric of a cognitive map is no longer adequate, EMs define instead an implicit universal value function that explains action selection by the ECM. But surprisingly, the ECM does not have to compute this universal value function: Its online learnt synaptic connections from indices for EMs to neurons that trigger actions automatically make it follow the gradient of this value function to the currently given goal, see our analyses in Section 2.5 and Section 4.6.2.

In contrast to the ECM algorithm, SR-based RL methods require computation and storage of a universal value function. This requires computing and storing of probabilities with high numerical precision. Storage of these probabilities in conductances of energy-efficient memristor arrays in edge devices is also problematic if they require high bit precision. It is actually also not plausible that synapses in the brain store these probabilities in their weights, because experimental data from neuroscience (Bartol Jr et al., 2015; Samavat et al., 2024) suggest that biological synaptic weights have a resolution of less than 5 bits. The ECM algorithm, which requires in its basic form only binary weights, is consistent with these constraints. In addition, we have shown in Fig. 6 A, B that the computational effort which the ECM algorithm requires for learning navigation in MDPs is substantially smaller than that for SR-based RL. Apart from learning, also inference, i.e., action selection, is substantially more efficient for the ECM algorithm: Unlike the SR, it does not have to compute value estimates for all possibly resulting next states, compare them, and then choose the one with maximal value estimate. This substantially reduces its latency for action selection. Apart from these advantages in the computational complexity of learning and inference, the ECM also has over SR-based RL the advantage that it automatically creates a cognitive maps that provides a “sense of direction” for navigation to any given goal in environments where actions are reversible. From the mathematical perspective, this reversibility condition is actually necessary for the existence of a cognitive map, since the distance measures in a cognitive map is automatically symmetric (according to the mathematical definition of a distance metric). But such a symmetric distance measure in a cognitive map is inconsistent with non-reversible actions, since this causes in general differences in the required effort to get from A to B versus getting from B to A. In addition, the ECM algorithm offers fast context adaptation and explainability, for which SR-based RL has no provision.

Our work makes a number of predictions for goal-directed action selection in the brain that can be tested through biological experiments. One prediction is that the fixation of a goal activates index cells for EMs that contain this goal. Furthermore, the action-selection mechanism of the ECM predicts that, at least when action selection has to be fast, the brain does not have to consider all possible options for action selection and call up for each of them the expected value. Finally, it predicts that index cells for EMs are able to activate neurons that can trigger actions which occured in the corresponding episode. There exist already experimental data which qualitatively support such direct links from EMs to actions (Robinson et al., 2020; Kim et al., 2023; Boscheron et al., 2026), but further experiments have to make the underlying synaptic connections and their plasticity explicit.

A limitation of this paper is that we only applied the ECM algorithm to tasks where states were uniquely labeled. In current work we extend this approach to tasks where the agent receives instead high-dimensional sensory inputs from the environment, and has to learn which features matter for getting rewards, see e.g. the discussion in (Gershman and Daw, 2017). Another challenge will be to extend our approach to non-Markovian situations, such as POMDPs (partially observable Markov Decision Processes), where the current sensory input does in general not provide enough information for inferring the current state. An obvious extension would be to activate index cells in a graded manner based not only on a given goal state and the current state, but also on recently encountered states, including states that are not identical to previously encountered ones. We conjecture that the action selection strategy of the ECM algorithm can be extended to these cases. We also did not try to apply clever exploration strategies or learning of affordances of actions in this version, but it should be easy to add them.

Further work will also examine a simplified version of the ECM algorithm (which we call GCM algorithm) that does not require index cells for episodic memories. This variant is based on the observation that it suffices for the goal-directed action selection of the ECM algorithm if not index cells, but neurons that represent states are linked to neurons that are able to trigger actions. The GCM algorithm is equally good at navigation to arbitrary goals in an MDP^-R^, but does not provide explainability or ad hoc adjustment to new contingencies. We propose that the GCM algorithm provides a suitable basis for modeling goal-directed action selection in the rodent brain, in particular for explaining the experimental data of (Zutshi et al., 2025). It also appears to be suitable for autonomously learning edge devices with more severe resource limitations.

## 4 Methods

### 4.1 Mathematical description of the ECM algorithm for the case where actions are reversible

We consider a task space that is defined by some undirected graph with *N*_*s*_ nodes. These nodes represent states *s*, which may include external and internal signals in real-world applications, or clusters/classifications of such signals that represent clusters of functionally equivalent observations, e.g. all observations made at a particular spatial location. For simplicity, we represent each state by a 1-hot code with an *N*_*s*_-dimensional binary vector. One could just as well use sparse binary codes, as long as these are approximately orthogonal (see Section A of the Supplement). There exists substantial evidence that orthogonal representations of states are prevalent in higher brain areas (Sun et al., 2025).Then for each EM *e* a linear neuron with a threshold, i.e., a perceptron, can be selected randomly to serve as index cell *z*_*e*_ for *e*.

#### Learning

Background from biology: Experimental data on synaptic plasticity in area CA1 of the rodent brain suggest that BTSP (Behavioral Time scale Synaptic Plasticity), creates index cells in the hippocampus (Bittner et al., 2017). Note that an application of BTSP during the commonly observed 10 to 20-fold speedup in replays of episodic memories (Takahashi, 2015; Michelmann et al., 2019) can substantially increase the temporal stretch of an episode to which an index cell responds. A long time window for synaptic plasticity has more recently also been observed in neurons on layer 5 of the rodent neocortex (Yaeger et al., 2025; Xiao et al., 2025). Hence it is reasonable to assume that a similar form of synaptic plasticity creates index cells for EMs also in the human brain. We use the simple model for BTSP from (Wu and Maass, 2025) that was shown to capture -in spite of its simplicity- the essence of biological data on BTSP.

We assume that *N*_*e*_ exploration episodes have been carried out in the task space. Each episode consists of a sequence of states, with every pair of consecutive nodes connected by an edge of the graph. For each episode *e*, we create an index neuron *z*_*e*_. A one-shot application of the simplified BTSP rule from (Wu and Maass, 2025) strengthens the synapse from every state neuron that is active during episode *e* to *z*_*e*_. The resulting input weights make *z*_*e*_ an OR gate over all states visited in that episode. For a state represented by a one-hot vector **s**, the activity of the index neuron is

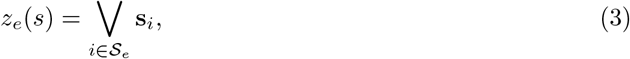

where *S*_*e*_ denotes the set of states visited in episode *e*, and V denotes the logical OR operation. Thus, *z*_*e*_ becomes active whenever the current state occurs in episode *e*. In the brain, several neurons may fire across all phases of an episode, but the ECM algorithm requires only one such index neuron for each episode.

The input weights of the *N*_*e*_ index neurons together form the embedding matrix 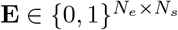. Each row of **E** contains the input weights of one index neuron *z*_*e*_. Assuming one-hot encoding of states, with **E**_*e,i*_ = 1 when state *i* occurs in episode *e*. We then use **E** to embed the *N*_*s*_ nodes of the graph into the *N*_*e*_-dimensional embedding space created by the episode-specific neurons. For a state *s* encoded by a one-hot vector **s**, the product **Es** activates exactly those index neurons whose episodes contain *s*.

Each action *a* is represented by an action neuron *n*_*a*_, whose activation can trigger execution of that action. The incoming weights of *n*_*a*_ mirror the embedding of the state reached after executing *a*. Let 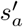 denote the resulting state. The incoming weight vector of *n*_*a*_ is

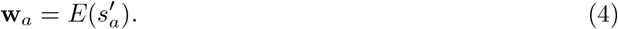

Thus, neuron *n*_*a*_ receives input from exactly those index neurons whose episodes contain the resulting state of action *a*. Collecting the incoming weight vectors of all *N*_*a*_ action neurons as rows forms the matrix 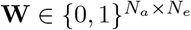.

#### Action selection

When producing an action sequence to reach the target state *g*, the goal embedding *E*(*g*) activates the action neurons through their incoming weights **w**_*a*_. Actions that cannot be executed in the current state are assumed to be inhibited (Stöckl et al., 2024). A WTA competition among the remaining action neurons then selects the most strongly activated action, as described in the main text, Eq. (1).

#### Alternative EM-based action selection

An alternative method for goal directed action selection is possible when each EM stores not only the sequence of visited nodes, but also the actions that were executed along the trajectory. In this case, action selection does not require learning synaptic weights from a representation of the next state after an action, as shown in Fig. 1E. Instead, the synaptic weight from the index neuron of an EM *e* to an action neuron *n*_*a*_ can be set to 1 whenever action *a* occurred in the corresponding trajectory. In the brain, this could correspond to learning backward connections from index neurons in CA1 to neocortical neurons that encode the content of the corresponding EM. Such learning is assumed to occur during replay (Chen and Wilson, 2023). Goal directed action selection can then be carried out through these learned synaptic weights using the same mechanism as shown in Fig. 1F.

### 4.2 Context-dependent selection of episodic memories

The basic ECM algorithm can be extended to context-dependent planning by gating episodic memories on or off. Each episode is tagged with the context in which it was stored. During planning, episodes that match the current context remain active, while the others are suppressed. The stored episodes and action weights remain unchanged.

This mechanism is used in the context-dependent planning experiments in Fig. 4. In these experiments, changing the context changes only which episodes are allowed to contribute to action selection.

Let *E*(*g*) denote the episodes that contain the current goal, and let 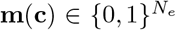 denote the context-dependent gate. The episodes available for planning are

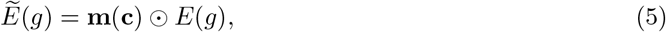

where *⊙* denotes elementwise multiplication. The input to each action neuron is then 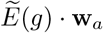. Thus, context adaptation consists simply of changing which stored episodes can vote for an action. The selected action can still be traced to the active episodes and their context tags.

### 4.3 Visualization of cognitive maps for graphs with invertible actions

For visualization in Figs. 2 and 4, the high-dimensional node embeddings were projected onto their first three principal components using PCA (Jolliffe, 2011). The resulting vectors were normalized to unit length and plotted on the surface of a sphere. Nodes with similar episode-membership patterns therefore appear closer together. This projection is used solely for visualization and does not affect learning or action selection.

### 4.4 Details to action selection in general graphs in Fig. 2

We created random instances of deterministic graphs by specifying the number of nodes and a connection probability between any pair of nodes. The edge probability was set to 3% for the 100-node graph.

For each deterministic graph, the ECM algorithm learns the environment from exploration episodes. We uniformly sample an equal number of random walks of length *L* = 16 starting from each node. For example, in a 100-node graph, sampling 10 trajectories per node results in *N*_*e*_ = 100 *×* 10 = 1000 episodes. At each step of a trajectory, the agent randomly selects one of the current node’s outgoing edges with equal probability and moves to the next node. A trajectory of length *L* contains *L* edges, corresponding to *L* + 1 nodes. Some nodes may appear more than once within an episode; however, we only record whether a node occurs (1) or not (0). Multiple occurrences of the same node within a trajectory do not affect its embedding.

As a baseline comparison, we employ the Dijkstra graph search algorithm. Dijkstra’s algorithm is an offline action selection method that requires unrolling the entire solution path before executing the first move. It generates the shortest path between any given start node and goal node.

Both matrices **E** and **W** are initialized to zeros. During action selection, an affordance gating mechanism is applied prior to the WTA process to inhibit actions that cannot be executed (see (Stöckl et al., 2024) for details).

#### 4.4.1 Graded ECM

To model the case where episodic neurons respond more strongly to nodes occurring near the center of a trajectory, we introduce a graded activation profile along each episode. For an episode *e* of length *L*, let *t ∈ {*0, …, *L}* denote the temporal index of a node within the episode. The activation weight *g*(*t*) assigned to that node is defined by a Gaussian function centered at the middle of the trajectory:

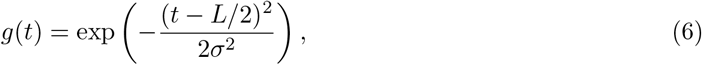

where the standard deviation *σ* controls the width of the bell shape. In our implementation, we set 4*σ* = *L*, so that the Gaussian effectively spans the entire episode length, and automatically adjusts when the trajectory length *L* updates.

Each node occurrence within an episode contributes to its embedding in **E** proportionally to *g*(*t*) rather than as a binary indicator:

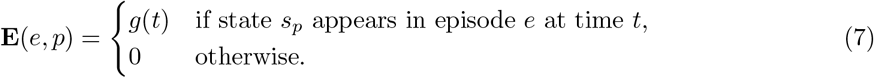

Thus, nodes near the middle of an episode receive higher activation values, while nodes at the beginning or end contribute less. If a state appears several times in an episode, we assign to it the highest value among all appearances. This graded mapping leads to a continuous-valued **E** matrix and improves action selection performance when only a limited number of exploration episodes are available (see Fig. 2C), and when the trajectory length is long (see Fig. 2D).

### 4.5 Details to the ECM algorithm with recursive self-improvement (RSI) in Fig. 3

For recursive self-improvement, the ECM algorithm keeps a fixed size pool of EMs and updates only its content. At each training step, a start and goal pair is sampled, and the current memory pool is used to generate a trajectory. ECM algorithm tends to reuse transitions that have frequently appeared on trajectories leading to the goal. This can reinforce poor early trajectories. We therefore add Gaussian noise to the inputs of the action neurons before the WTA operation during training. The noise is scaled by the current action-neuron activations and decreases linearly from 0.35 to 0.05. No such training noise is used during evaluation. The input **I**_*t*_ to all action neurons then becomes:

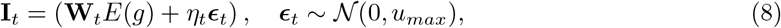

with

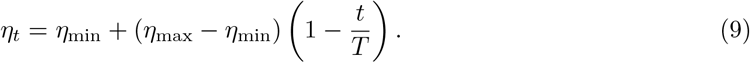

Before a generated trajectory is written into memory, loops are erased by removing segments between repeated visits to the same node. The cleaned trajectory is then relabeled by its actual endpoint, following the idea of Hindsight Experience Replay (Andrychowicz et al., 2017). We also consider all subpaths of the cleaned trajectory as candidate EMs, with each subpath labeled by its own start and endpoint.

The brain has complex mechanisms for deciding which experiences should be retained, forgotten, or compressed into a more general schema. In our implementation, we approximate this memory selection process by assigning a quality score to each stored episode. Each candidate EM is scored by how many ordered start and goal pairs it adds to the memory pool, or how many existing pairs it improves by providing a shorter stored route. This contribution is normalized by trajectory length, with a small length bonus of *ρ* = 0.20 to avoid storing only very short fragments. When the pool is full, a candidate replaces the currently weakest EM only if its score is higher.

For online methods, convergence also depends on which tasks are practiced. It is therefore useful to sample start and goal pairs actively from relations that are less explored or currently difficult to solve. We sampled training pairs from a mixture of random pairs and hard pairs, with probability 0.5 for each source. Hard pairs are pairs for which the current memory has no stored route, fails to reach the goal, or produces a route longer than 1.5 times the best stored route for the same pair.

### 4.6 Markov Decision Processes without reward functions (MDP^-R^s)

Navigation in Markov decision processes (MDPs) is a standard benchmark task in RL. Since action outcomes are here stochastic, one needs to specify a policy, rather than just a path for reaching a goal node from a given start node. But most RL algorithms consider only the case where the goal (reward function) is fixed. In contrast, the ECM algorithm produces, after exploring the structure, a policy for reaching any given goal. Since there is apparently no standard notation for MDPs without a reward function, we refer to them in this work as MDP^-R^s. Such structures describe only the stochastic state transitions that are caused by actions, and serve as platforms on which different goal-directed behaviors can later be planned when given a goal.

Each MDP^-R^ consists of *N*_*s*_ nodes (states) and *A* actions per node, resulting in *N*_*a*_ = *N*_*s*_ *× A* distinct actions.

Each state *s* is represented by a one-hot vector 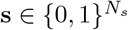. Similarly, each action *a* is also defined by a one-hot action vector 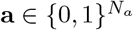 with the *a*^th^ element set to 1 and all others are set to 0s.

We created generic MDP^-R^s for our experiments using the widely adopted Garnet procedure to generate large MDPs (Bhatnagar et al., 2009). In our implementation, each state had either two or three available action commands, corresponding to panels in Fig. 5C and Fig. 6. We added a directed cycle as one of the action outcomes of one of the actions through all states to ensure communication. Transition probabilities were sampled from a Dirichlet distribution with *α* = 0.8, and self-loops were not allowed.

These definitions specify the stochastic environment used by all algorithms in the study. Since no reward function is assigned, the same learned transition structure can be reused for action selection toward any desired goal.

#### 4.6.1 Mathematical description of the ECM algorithm for MDP^-R^s

For each episode *i*, we update the state embedding matrix 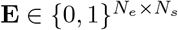 by the same rule as Eq. (3) as for deterministic graphs.

The 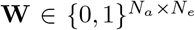 matrix is updated by having a synaptic connection from a neuron *z*_*e*_ in CA1 that represents an EM *e* to an action neuron *n*_*a*_ only if *z*_*e*_ is the index for an EM *e* that starts with this action *a*:

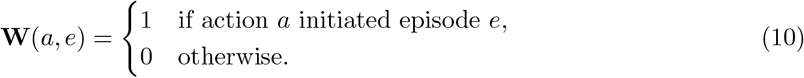

The action selection mechanism remains the same as in the deterministic case. The goal embedding *E*(*g*) activates all episodic neurons *z*_*e*_ whose corresponding episodes contain the node *g*. Through the incoming weights **w**_*a*_, these neurons activate each action neuron *n*_*a*_ according to how often episodes initiated by action *a* reached the goal state. A WTA mechanism applied to the action neurons then selects the most strongly activated action.

#### 4.6.2 Universal value function underlying ECM action selection

For MDP^-R^s, the ECM algorithm chooses that action for which exploration episodes initiated by this action reached the goal most often. Here we show that, under uniform random exploration, this comparison of episode counts asymptotically implements a greedy step with regard to a universal value function.

Let *C*_*L*_(*s, g*) count exploration episodes of length *L* starting from *s* that contain *g*, and let *N* (*s*) count all episodes starting from *s*. We define the universal value function as

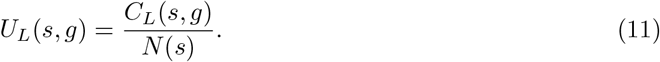

This estimates the probability of reaching *g* within *L* transitions under the exploration policy. The initial state is included, so *U*_*L*_(*g, g*) = 1. The corresponding value *U*_*L−*1_(*s, g*) counts goal visits within the first *L −* 1 transitions.

Greedy action selection compares the expected values *U*_*L−*1_(*s*^*′*^, *g*) of resulting next states *s*^*′*^. These values are estimated from episodes starting at *s*^*′*^, whereas ECM uses only episodes initiated by the candidate action *a* in state *s*. We therefore need to establish that the continuations of these action-specific episodes lead to the same estimates.

For *s ≠ g*, let *N* (*s, a, s*^*′*^) count episodes whose first transition is 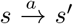, and let *C*^*L−*1^ (*s, a, s*^*′*^, *g*) count those whose continuation, including *s* and the remaining *L −* 1 transitions, contains *g*. The Markov property ensures that the probability of reaching *g* within these remaining transitions is independent of how *s*^*′*^ was reached, provided subsequent actions follow the same exploration policy. Hence,

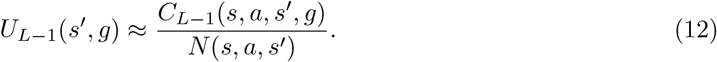

The approximation reflects finite sampling, with its error vanishing as exploration increases.

Let *N* (*s, a*) count all episodes starting from *s* with action *a*, and let *C*_*L*_(*s, a, g*) count those that contain *g*. Substituting the empirical transition probability 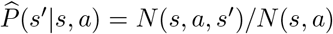 and Eq. (12) into the expected next-state value gives

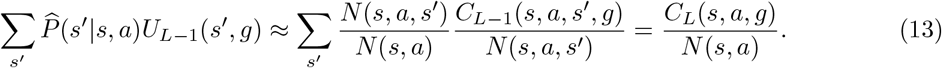

The sums run over sampled successor states. The equality follows by cancelling *N* (*s, a, s*^*′*^) and summing successful episodes over their first successor states.

Random-walk exploration makes the denominator in Eq. (13), *N* (*s, a*), approximately equal across available actions, so comparing *C*_*L*_(*s, a, g*) gives the same action ranking as comparing the normalized values. By Eq. (10), *E*(*g*) *·* **w**_*a*_ = *C*_*L*_(*s, a, g*). Thus, the WTA mechanism asymptotically implements a greedy step with regard to *U*_*L−*1_(*s*^*′*^, *g*), without explicitly computing successor-state values.

### 4.7 Details to MDP^-R^s in Fig. 5 and Fig. 6

We created random MDP^-R^s with *N*_*s*_ nodes, where *N*_*s*_ is specified for each experiment. We used *A* = 2 actions per node in Fig. 5 and Fig. S2, and *A* = 3 in Fig. 6. Each state-action pair had *K* = 2 possible successor states.

Episodes were generated by random walk exploration in the MDP. Each episode starts from a uniformly sampled node *s*^0^ *∈ {* 1, …, *N*_*s*_*}* and runs for a fixed length *L*. At each time step *t*, an action *a*^*t*^ *∈ {*1, …, *A}* is chosen uniformly at random, and the next state *s*^*t*+1^ is drawn from *P* ( *·*|*s*^*t*^, *a*^*t*^). This procedure ensures that, in expectation, each action is applied equally often at every node, introducing no sampling bias when later comparing which action most frequently leads the agent toward the goal.

#### 4.7.1 Implementation of RL algorithms for comparison

##### Value iteration

Value iteration provides a gold standard for navigation in MDPs since it is guaranteed to converge to an optimal policy (Sutton and Barto, 2018). However, it needs to be rerun for each goal node.

Our implementation follows the value-iteration algorithm in (Sutton and Barto, 2018), Section 4.4 (“Value Iteration”). We use its equivalent unit-cost formulation for stochastic shortest-path problems:

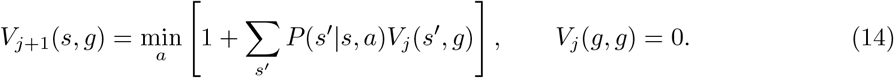

Each transition incurs a unit cost of 1 until the goal is reached, and the goal state has value *V* (*g, g*) = 0. Thus, *V* (*s, g*) represents the expected number of transitions required to reach goal *g* under the current value estimate. The algorithm loops over all states for every possible goal, assuming that the complete transition probability matrix **P** is known in advance. The greedy policy is extracted at the evaluation checkpoints specified in Sec. 4.7.6 as:

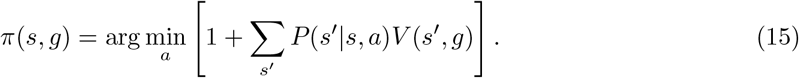

A separate value function is required for each goal. In our benchmark, value iteration is given the exact transition matrix **P** in advance, whereas SR, Q-learning, and ECM learn from sampled experience without access to this pre-specified transition model. We therefore count the cumulative VI cost required to compute value functions for all possible goals.

##### Successor representation

For the Successor Representation (SR) baseline, we followed the original temporal-difference formulation introduced by (Dayan, 1993). The SR matrix **M** encodes the expected discounted future state

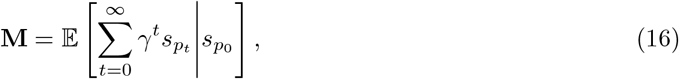

where *p*_*t*_ denotes the index of the state visited at time *t*, and 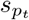 is its corresponding one-hot vector. Thus, each row **M**(*s*_*p*_) represents the expected discounted visitation frequencies of all future states when starting from state *p*. The matrix satisfies the fixed-point relation **M** = **I** + *γP*_*π*_**M**, where *P*_*π*_ is the transition matrix under policy *π*.

We implemented the standard on-line one-step temporal-difference (TD) learning rule:

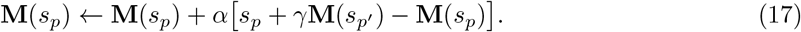

where *s*_*p*_ and 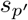 correspond to the current and next states, respectively. We used a learning rate of *α* = 0.1 and a discount factor of *γ* = 0.95. The behavior policy during training was uniformly random over actions to ensure sufficient exploration.

During exploration, the same observed transitions were also used to estimate the action-conditioned transition probabilities 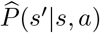. For a given goal state *g*, the value of selecting action *a* in state *s* was computed as

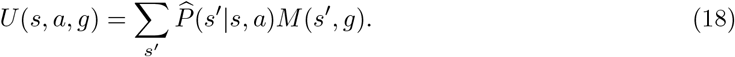

The corresponding goal-conditioned policy was

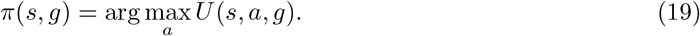

The same learned SR matrix can be reused for different goals by selecting the corresponding goal column *M* (:, *g*).

##### Q-learning

We implemented tabular goal-conditioned Q-learning in a unit-cost formulation with all-goals updating, closely related to the DG-learning formulation of Kaelbling (1993), using standard temporal-difference updates (Sutton and Barto, 2018). The table *Q*(*s, a, g*) stores the estimated number of remaining steps after selecting action *a* in state *s* while pursuing goal *g*. During exploration, actions were selected uniformly at random. Each observed transition (*s, a, s*^*′*^) was shared across all possible goals, and the Q-values were updated according to

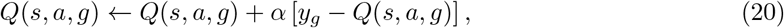

where

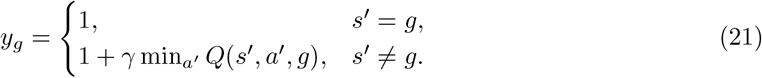

We used *α* = 0.1 and *γ* = 1. The corresponding goal-conditioned policy was

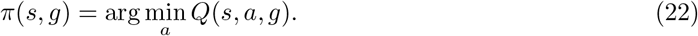

Sharing a transition across all goals reduces the required number of interactions with the environment, although up to *N*_*s*_ goal-conditioned Q-values are still updated after each transition.

#### 4.7.2 Measures for evaluating the computational complexity of four approaches in Fig. 5C and Fig. 6A-B

The multiply–accumulate (MAC) operation is a dominant arithmetic operation in many neural-network workloads (Sze et al., 2020). It updates an accumulator according to *c ← a × b* + *c* and is conventionally counted as one multiplication and one addition, corresponding to two floating-point operations (FLOPs). Under a simple accounting scheme without operand reuse, register retention, or caching, each MAC also involves three reads for *a, b*, and *c*, followed by one write of the updated value of *c*. This corresponds to four algorithm-level memory accesses. Actual physical memory traffic depends on the hardware architecture and data-reuse strategy and therefore cannot be inferred directly from the MAC count. Moreover, algorithms can perform memory reads, writes, and logical operations without executing any floating-point arithmetic, as in ECM. We therefore report MACs and FLOPs separately.

The estimates include operations required to update the learned representations. For ECM, SR, and Q-learning, random-walk trajectories are treated as observations from the environment, and the operations used to generate these trajectories are not included. Policy evaluation at the reported checkpoints is also excluded, as are comparisons for action selection, array indexing, and loop-control operations. The reported costs therefore quantify algorithm-level learning updates rather than the instructions or runtime of the software implementation used to simulate them. For VI, the transition model is provided in advance, and repeated memory accesses to its fixed transition probabilities are not charged. Values that can remain locally in an accumulator or register are not counted as additional memory accesses.

The complexity of learning is of a somewhat different nature for the four algorithms that are compared in Fig. 5 and 6. VI is given the complete transition model, but requires repeated floating-point Bellman backups and a separate value map for each goal. Q-learning learns from experience, but repeatedly updates goal-dependent action values, even though each sampled transition is shared across all goals in our implementation. Like the ECM algorithm, SR can reuse a learned representation when the goal changes. Its successor matrix, however, contains floating-point weights that require repeated updates and describe a fixed behavioral context. The ECM algorithm stores binary weights, each of which is updated at most once. It can reuse the same episodic memories for new goals and adapt immediately to a new context by selecting the memories relevant to the current situation.

##### Value iteration

For each goal, one value-iteration sweep updates every state according to

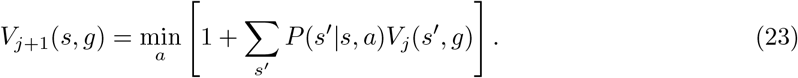

This is the unit-cost formulation used in our implementation. Each state–action pair has on average *k* possible successor states, with *k* = 2 in our experiments. For each successor, multiplying *P* (*s*^*′*^ | *s, a*) by *V*_*j*_(*s*^*′*^, *g*) and adding the result to the accumulator requires one MAC, corresponding to two FLOPs. The unit transition cost is incorporated by initializing the accumulator to 1 and is therefore not charged as a separate arithmetic operation in our accounting.

The benchmark computes value functions for all *N*_*s*_ possible goals. One complete sweep therefore requires

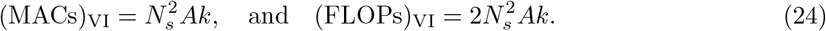

We assume in Fig. 5C that VI has exact access to the transition matrix **P** of the MDP^*−*R^. If VI also had to explore the environment, as SR, Q-learning, and ECM do, it would incur additional costs for collecting transitions and estimating **P**. For example, applying every action 10^3^ times at every state would require *N*_*s*_*A ×* 10^3^ observed transitions and corresponding updates to the transition counts. These updates would contribute additional memory accesses and arithmetic operations, depending on how the transition probabilities are estimated and normalized. A value function computed from an estimated transition matrix is also optimal only for the estimated model and may therefore differ from the optimal value function of the true MDP.

VI also cannot directly reuse its learned value function when the goal changes. Each goal defines a different reward or cost function, requiring VI to recompute the value table. By contrast, SR can reuse its learned successor matrix for different goals, while ECM can retrieve goal-relevant episodes without relearning its representations. We therefore report the cumulative cost of running VI separately for all possible goals.

##### Successor representation

For SR, every observed transition from *s* to *s*^*′*^ produces the update

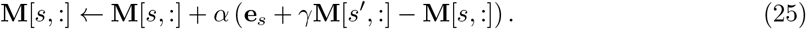

The update is applied to all *N*_*s*_ entries in one row of the successor matrix. For the MAC-based comparison, we approximate the update by two effective MACs per matrix entry. For the FLOP count, we exploit that **e**_*s*_ is one-hot. For all but one of the *N*_*s*_ entries, its contribution is zero, so the update requires approximately four floating-point operations per entry. The single additional operation for the non-zero entry of **e**_*s*_ is omitted from this dominant-term estimate. The resulting costs are

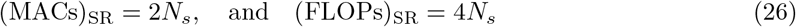

per transition. The single addition used to update the observed transition count is omitted from this dominant-term estimate.

For an episode containing *L* transitions, these two costs in Eq. (26) are multiplied by *L*.

SR in addition needs the transition counts because a state-based SR represents expected future state occupancy but does not directly specify which action leads to each successor state. The counts are used to estimate **P** when extracting a goal-conditioned policy. The normalization of these counts and the matrix multiplication used for policy extraction are performed only at evaluation checkpoints and are excluded from the reported learning costs.

Alternatively, SR can be constructed from transition counts after exploration. During learning, each observed transition requires only one integer count increment, with no floating-point MACs or FLOPs. After learning, the counts are normalized and averaged over actions to obtain the uniform-policy transition matrix 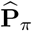, requiring *O*(*N*_*s*_*Ak*) operations with sparse storage. The successor matrix is then computed as 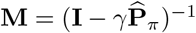. Standard dense LU-based inversion, including factorization, requires approximately 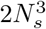 FLOPs, or 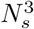 MAC-equivalents, to leading order. For *N*_*s*_ = 1000, this amounts to approximately 2 *×* 10^9^ FLOPs or 10^9^ MAC-equivalents per construction. This alternative replaces repeated TD updates with post-learning SR construction before planning. Its total computational cost is still on the order of 10^9^, offering no advantage over ECM when compared in Fig. 6. Our reported costs use TD-based SR; comparisons with this alternative should include its construction cost whenever the SR is recomputed.

##### Q-learning

The Q-learning implementation reuses every observed transition to update the values for all possible goals:

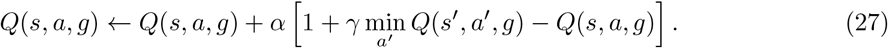

For each goal, computing 1 + *γ* min_*a*_*′ Q*(*s*^*′*^, *a*^*′*^, *g*) requires one MAC after the minimum has been found. Applying the temporal-difference correction requires one subtraction and a second MAC. The approximate arithmetic cost per transition is therefore

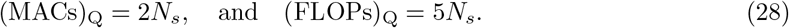

The comparisons used to find the minimum action value are not counted as FLOPs.

These expressions slightly overestimate the exact cost because the update for the current goal is skipped, while reaching the goal removes the need to read the next-state action values. For an episode of length *L*, all three costs are multiplied by *L*.

##### ECM

During learning, ECM updates one row of the episode matrix **E** for each exploration episode. The update sets binary entries indicating which states occurred in the episode and requires no floating-point arithmetic. ECM also updates one entry in the action matrix **W** to associate the episode with its initiating action according to Eq. (10).

For comparability with SR, Q-learning, and VI in the MAC-based analysis, we count each binary memory write performed by ECM as one effective MAC. This is a conservative accounting convention rather than a count of actual MAC operations. Under the simple accounting scheme used here, one MAC contains two FLOPs, three memory reads, and one memory write, while an ECM update only sets a binary weight. This convention therefore places ECM on the same scale as the arithmetic-based algorithms while overestimating its arithmetic cost.

An episode containing *L* transitions contains *L* + 1 visited-state occurrences. Updating the corresponding row of **E** therefore requires at most *L* + 1 binary writes, since repeated visits to the same state do not require additional entries to be set. Updating **W** requires one additional write. The effective MAC count is therefore approximated as

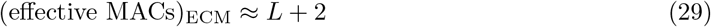

per episode. This value is added to the MAC counter after each episode during learning. The same updates require no floating-point operations:

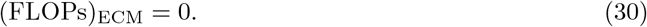

Logical operations used to identify visited states and set binary entries are not counted as FLOPs. Each binary weight needs to be set at most once and does not require repeated numerical adjustment.

#### 4.7.3 Graded index-cell activation based on episode safety

To incorporate safety into ECM-based path finding in Fig. 5F, each activated index cell is assigned a graded activation according to the overall safety of its corresponding episode. For an episode 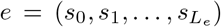, where each node *s* has risk probability *p*_risk_(*s*), episode safety is defined as 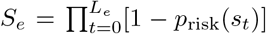. This safety score is then converted into an activation weight using *w*_*e*_ = exp[ *−β*(1 *− S*_*e*_)], where *β* = 25 controls the strength of safety modulation. Safer episodes therefore retain stronger activation, whereas less safe episodes are increasingly suppressed.

If *z*_*e*_ *∈ {*0, 1*}* denotes the original binary activation of index cell *e*, its graded activation becomes 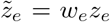. These graded activations are used directly for action voting, so safer retrieved episodes contribute more strongly to action selection. This modulation is applied only during retrieval and does not require relearning the stored episode matrix **E** or action-association matrix **W**.

#### 4.7.4 Safety modulation in 1000-node MDP^-R^s (Fig. 6C)

We evaluated 10 independently generated MDP^-R^s with 1000 states, three actions per state, and two possible successors per action. Each instance was explored using 10^6^ random-walk episodes of 20 transitions, with uniformly sampled starting states and actions. Each episode was associated with its initiating state–action pair and contributed one vote for each distinct goal state it contained.

We assigned a fixed accident probability *p*_risk_(*s*) to each node *s*. First, separate rollouts of the ECM policy without safety modulation were used to identify the most frequently visited node. This node was assigned an accident probability of 0.3% per visit. We then randomly selected 49 distinct nodes from the remaining 999 nodes, with each node equally likely to be selected. Each selected node was assigned an accident probability of 0.001% per visit. The other 950 nodes had zero accident probability. This assignment rule was selected on a separate development environment and then applied to the 10 test instances.

Similarly as in Sec. 4.7.3, episode safety and graded activations *w*_*e*_ of EMs were computed by

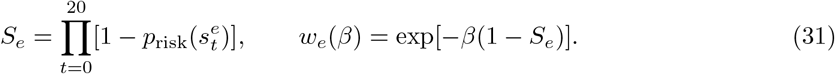

Repeated visits contributed separately to episode safety.

We evaluated *β ∈ {*0, 100, 300, 1000, 3000, 10^4^, 3 *×* 10^4^, 10^5^, 2 *×* 10^5^, 3 *×* 10^5^, 5 *×* 10^5^, 7.5 *×* 10^5^, 10^6^*}* while keeping memories, risk maps, and tie-breaking priorities fixed. At each navigation step, the action receiving the largest sum of weighted episode votes was selected.

Each setting was evaluated using one stochastic rollout for each of the 999,000 ordered pairs of distinct start and goal states. Rollouts ended at goal arrival or after 100 transitions; timeouts contributed a length of 100. For the visited sequence (*s*_0_, …, *s*_*τ*_ ), accident probability was 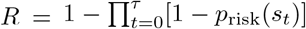, including starting states, reached goals, and repeated visits. Accidents were scored analytically without terminating navigation; unfinished rollouts contributed their observed-prefix risk. All pairs received equal weight, and transition random numbers were shared across *β* values. Curves and shading show means and *±*1 sample standard deviation across the 10 instances.

#### 4.7.5 Removal of episodic support for action selection (Fig. 6D)

ECM agents were trained on 10^7^ random-walk episodes of 20 transitions in each of 10 independently generated 1000-state MDP^-R^s, using the exploration and memory conventions above. Episodes contributed equal votes (*β* = 0). For each ordered pair of distinct start and goal states, we identified the action selected by the complete memory bank and its *m* supporting episodes. We removed *⌊pm/*100 *⌋*supporting votes, varying *p* from 0 to 100 in increments of 0.1, while preserving competing-action votes and tie-breaking priorities.

The intervention was applied independently to each pair and affected only the first action. All subsequent decisions, including revisits to the starting state, used the original full-memory policy. We measured mean navigation length across all 999,000 pairs. Evaluation followed the same one-rollout-per-pair protocol with a 100-transition cap, timeouts contributing 100, and shared transition random numbers across removal levels. Curves and shading show means and *±* 1 sample standard deviation across the 10 instances.

#### 4.7.6 Other detailed settings for performance comparison in Fig. 5C and Fig. 6

Performance was evaluated at fixed checkpoints during learning. For VI, a checkpoint is defined by the number of complete Bellman sweeps over all states and all possible goals. We evaluated VI after sweeps 1–20, and subsequently after 30, 50, 100, and 200 sweeps. Thus, the early part of the learning curve was sampled more densely, where performance changes most rapidly.

For ECM, SR, and Q-learning, a checkpoint is instead defined by the cumulative number of random-walk exploration episodes that have been processed. We used approximately logarithmically spaced checkpoints at 10, 20, 50, 100, 200, 500, 10^3^, 2 *×* 10^3^, 5 *×* 10^3^, 10^4^, 2 *×* 10^4^, 5 *×* 10^4^, 10^5^, 2 *×* 10^5^, 5 *×* 10^5^, 10^6^, 2 *×*10^6^, 5 *×* 10^6^, and 10^7^ episodes, truncated when the performance has converged. At each checkpoint, learning was temporarily stopped and the current policy was extracted. Performance was evaluated by one stochastic rollout for each ordered pair of distinct start and goal states, with a maximum horizon of 100 steps. The resulting path lengths were averaged across all start-goal pairs. Results in Figs. 5C and 6 were then averaged across 10 independently generated MDP^-R^ instances. The computations required for policy extraction and evaluation were not included in the reported learning costs.

The horizontal position of every data point in the computational-cost plots was obtained by converting the amount of learning completed at that checkpoint into the corresponding cumulative MAC, memory-access, or FLOP count using the accounting described above. Hence, the curves compare policy performance after progressively larger amounts of learning rather than after equal numbers of iterations or environmental observations across algorithms.

## 5 Acknowledgments

We would like to thank Christoph Stoeckl for inspiring discussions. This research was partially supported by the National Science Foundation of the USA (EFRI BRAID project 2318152) and the Austrian Science Fund (FWF) (10.55776/COE12). The authors also gratefully acknowledge the Gauss Centre for Supercomputing e.V. (www.gauss-centre.eu) for funding this project by providing computing time on the GCS Supercomputer JUWELS[1] at Jülich Supercomputing Centre (JSC). For open access purposes, the author has applied a CC BY public copyright license to any author accepted manuscript version arising from this submission.

## Supplementary Information

### A Non-orthogonal states

In abstract graph experiments, each node is encoded as a one-hot state vector, such that all states are orthogonal with each other. However, this is not a strict requirement for the ECM algorithm. In this section, we explain how our model can tolerate non-orthogonal inputs.

#### A.1 Non-orthogonal code generation

On the 32-node abstract graph environment, we generated for each node a 10^4^-dimensional binary vector with sparsity = 0.5% following the data 0.2% to 1% measured in the human medial temporal lobe (Waydo et al., 2006; Fried, 2022).

The degree of orthogonality is adjustable: The indices of ones are adjusted either to overlap with the dimensions of other items’ ones, increasing the average cosine similarity, or to shift overlapping ones to different dimensions, reducing the cosine similarity. For a given target average cosine similarity, this process is iterated for a sufficiently long time to achieve a numerically optimal coding configuration.

#### A.2 Training

The training still follows the same rule, except that the inputs are no longer one-hot vectors.

A trajectory contains *L* transitions, or *L* + 1 states. All index neurons that fire within a trajectory will induce a LTP on the incoming weights of a index neuron (with index *i*) that is selected by the plateau potential:

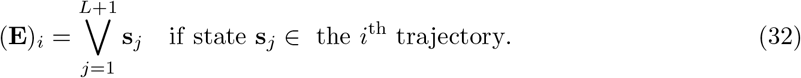

In above, the V represents dimension-wise “OR” operation between binary vectors: the result on a dimension is one if any state **s**_*j*_ has a one on this dimension.

#### A.3 State embedding

Non-orthogonality poses a challenge for our model. When a state lies outside of a trajectory but shares some “1” entries with an in-trajectory state, the off-trajectory node still activates the corresponding index neuron—and that neuron ideally shouldn’t respond. This undesired activation injects noise into the model.

Crucially, if only a subset of a state’s ones overlaps with another, then the index neurons encoding the true trajectory state should exhibit higher activations than those encoding the spurious overlap. To enforce this, we introduce a nonlinearity *f* (*·*) and replace the original state-embedding formula *E*(*s*) = **Es** with:

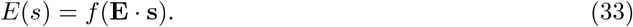

Specifically, we use a max-thresholding function that preserves only the maximal entries (zeroing out all others). This effectively suppresses the unwanted activations and eliminates noise. One can implement this also with a suitable global inhibition.

#### A.4 Result

We performed a control experiment using sparse, non-orthogonal state codes with the dimensionality and sparsity specified above. With the modified state-embedding procedure, ECM maintained near-optimal navigation performance. With the average pairwise cosine between states = 0.2 (highly non-orthogonal as compared with data from human MTL in 0.02 to 0.08 (Yang and Maass, 2026)), and trained on *N*_*e*_ = 1000 trajectories with each contains *L* = 10 nodes. The ECM agent on average takes 3.114 steps on a 32-node random graph, only 1.81% worse than Dijkstra’s 3.058 steps.

### B Navigation to arbitrary goals in directed graphs

Directed graphs are special cases of MDP^-R^s where the outcome probabilities of all actions are 0 or 1. In this case one can compute the shortest path via Dijkstra, like for undirected graphs.

**Fig. S1:**
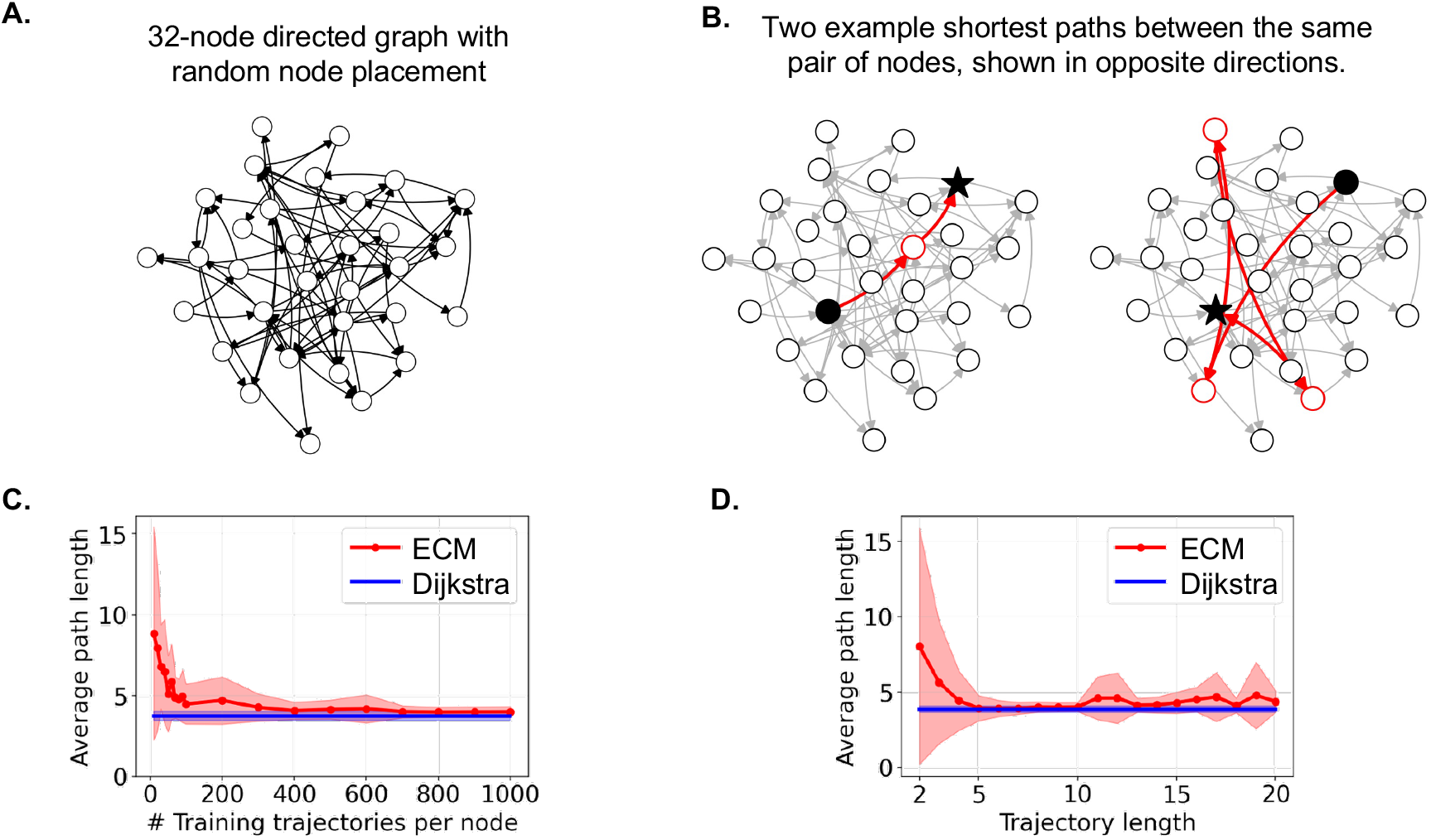
Performance of the ECM algorithm for goal-directed action selection in directed graphs. **A**. An example random directed graph. **B**. Two example shortest paths for a pair of nodes in opposite directions. The trajectories can be very different when switching start and goal nodes. **C**. and **D**. show mean *±* standard deviation over 10 randomly generated directed graphs. In C, one sees that the model already has near-optimal performance when the agent is trained on 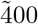 episodes per node (with trajectory length = 6). In D, one sees that the model has a very stable near-optimal performance for trajectory length ranging from 5 to 10 (with 1,000 trajectories per node).

Across 10 random directed graphs, our model’s average path length is **3.903**—only 4.51% longer than Dijkstra’s 3.734—using 1000 training trajectories per node (length 6).

### C MDP^-R^s

We provide two supplementary analyses of the experiment in Fig. 6, comparing VI, SR, Q-learning, and ECM on 1000-node MDP^-R^s with three actions per node.

Figure S2 complements the path-length comparison in the main text by reporting planning success rates as a function of the computational cost invested in learning. We quantify this cost using multiply–accumulate operations (MACs) and floating-point operations (FLOPs). Success rates measure the fraction of evaluation rollouts that reach the goal within the evaluation horizon of 100 steps. Results are averaged across 10 independently generated MDP^-R^ instances, with shaded regions indicating one standard deviation across instances (too small to be visible).

To assess the sensitivity of the performance estimates to stochastic action outcomes, we also repeat the evaluation using 10 rollouts for each start–goal pair, compared with one rollout per pair in Fig. 6. As shown in Fig. S3A, the resulting performance curves remain nearly unchanged. The standard deviation among the 10-rollout is shown in Fig. S3B. One sees that the value remains small. This agreement indicates that increasing the number of evaluation rollouts does not materially alter the aggregate comparison among the four algorithms in this setting.

**Fig. S2:**
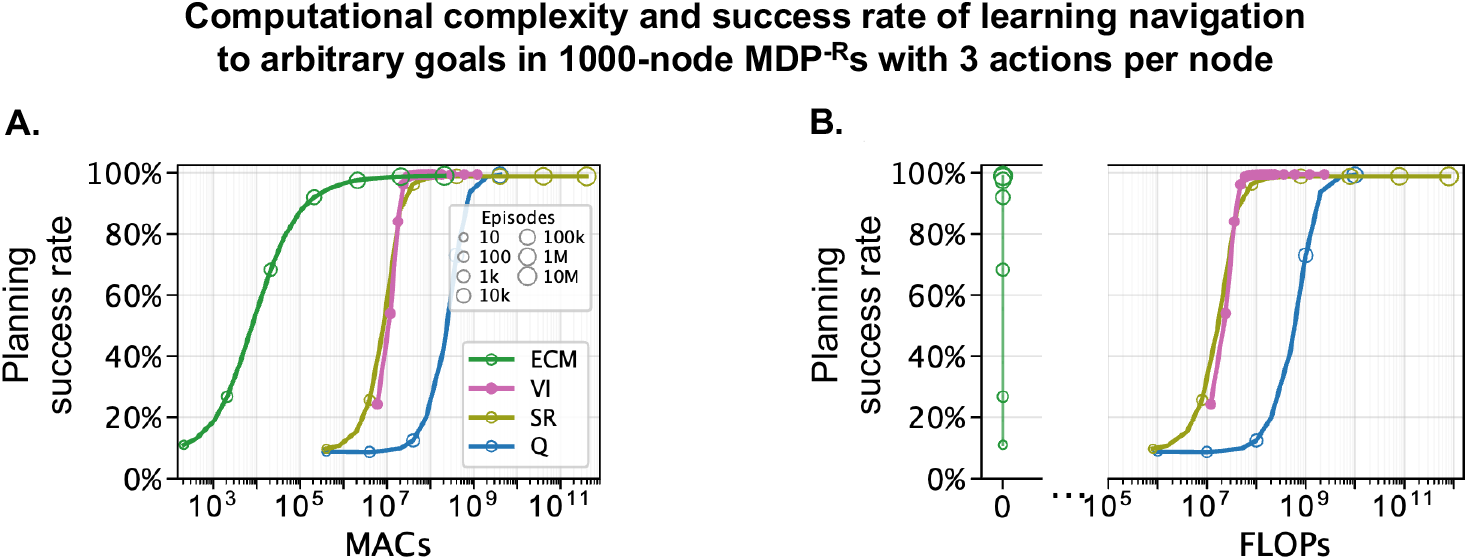
Success rate comparison of VI, SR, Q-learning and ECM on a 1000-node MDP^-R^, with three actions per node. **(A)** Average planning success rate as a function of the number of multiply–accumulate operations (MACs). **(B)** The same comparison as a function of the number of floating-point operations (FLOPs). Panels A–B show the mean *±* standard deviation (shaded) across 10 randomly generated MDP^R^ instances. The standard deviations are too small to be visible here.

**Fig. S3:**
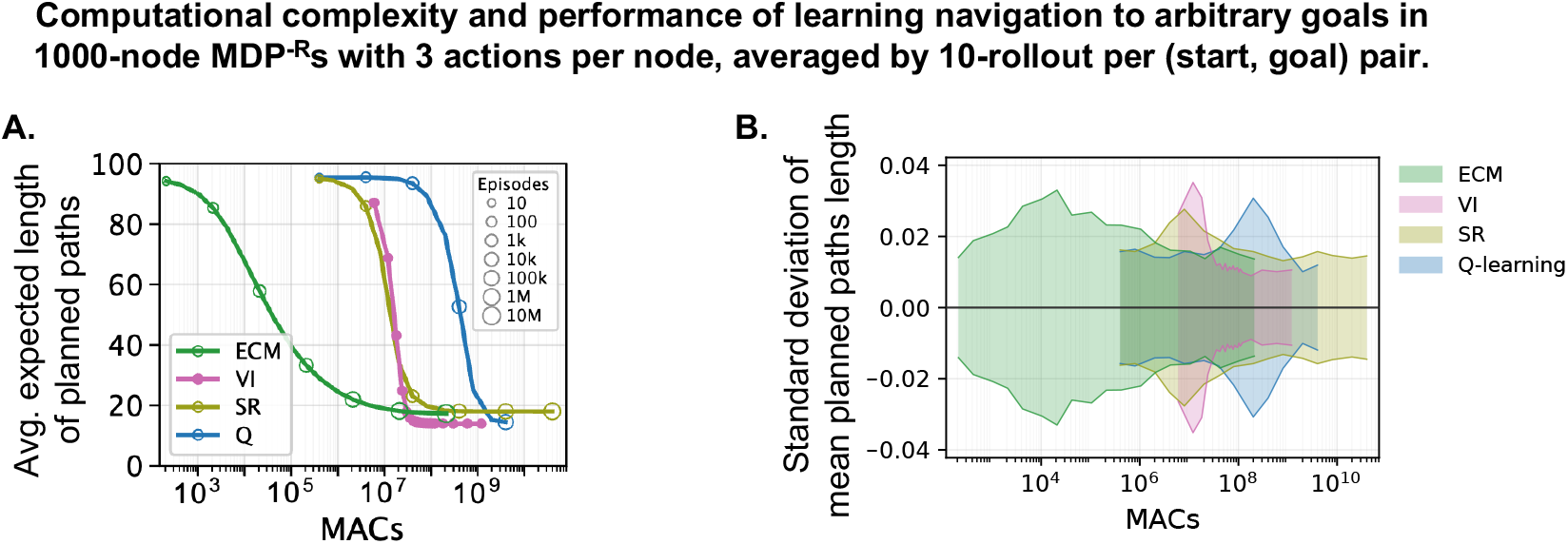
Comparison of VI, SR, Q-learning and ECM on a 1000-node MDP^-R^, with three actions per node and results averaged over 10 rollouts for each start and goal pair. **(A)** Performance is nearly identical to that in Figure 6, which uses a single rollout for each start and goal pair. **(B)** Shaded bands show *±*1 standard deviation of mean planned path length across 10 rollouts within each MDP, averaged over 10 MDP instances, as a function of learning MACs. Means are centered at zero to highlight rollout variability.

### D Foraging

We considered so far only applications of the ECM algorithm to tasks where reaching a particular state or location was specified as behavioral target. But the ECM-approach works just as well for other tasks, provided that the behavioral target can be defined in terms of a set *G* of EMs. The ECM algorithm generates then a sequence of actions that maximize correlation between the EM-based encoding of the resulting state and *G*.

**Fig. S4:**
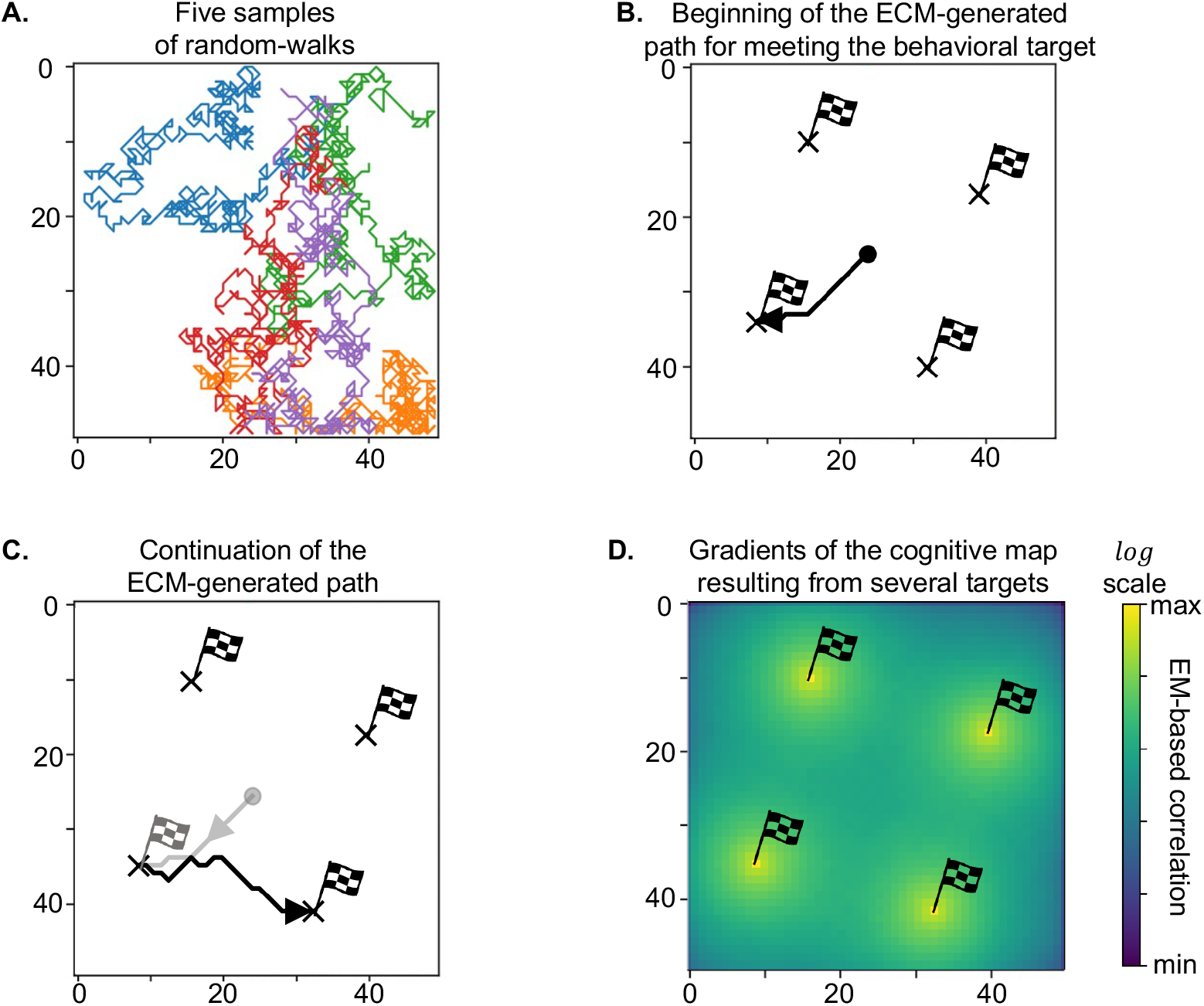
Application of the ECM algorithm to a foraging task. **(A)** 5 samples of random exploration trajectories, from random starting points. Each trajectory, consisting of 500 steps, gives rise to an EM. **(B)** Foraging from a start point in the middle between the 4 target points. The movement trajectory that is generated by the ECM algorithm immediately moves away from the center between them. **(C)** After reaching the first target point, the ECM algorithm automatically produces a trajectory to another one of them, once EMs that contain the first one are discarded (or the index neurons for them are inhibited). **(D)** Structure of the cognitive map that is produced by correlations between locations (states) and the set *G* of EMs that define the behavioral target. Locations between target points have low correlation with the target points in this cognitive map.

In particular, foraging tasks are supported, where a set of target points in a physical environment are given, and the target behavior is a short sequence of movements that visits each of them. This behavioral target is defined by the set G of all EMs where at least one of the target points was visited. An inherent algorithmic difficulty of value-function based RL solutions is that value functions arising from different target points superimpose at locations between target points, thereby attracting movements to these in-between locations.

We show in Fig. S4B that a straightforward application of the ECM algorithm, following a standard random exploration of the environment (see Fig. S4A) does not face this difficulty. It produces a trajectory that moves from the center in between the 4 target points to one of them. If one removes then those EMs from G that had visited this point, the continuation of the trajectory that is generated by the ECM algorithm moves on to one of the other target points. Hence it generates a meaningful foraging behavior.

The reason why the ECM algorithm does not fall into the trap of moving towards locations between several target points is illustrated in Fig. S4D: The EM-based correlation of these locations (states) with the behavioral target that is defined by G is fairly low at these in-between locations. This results from a fast decay of the probability that an exploration trajectory that visits one of the 4 target points also visits a specific location in between them when the distance of this location to the nearest target point becomes larger.

#### D.1 Further details to foraging (multi-goal navigation) in a 2D discrete environment

We evaluated the agent in a two-dimensional discrete navigation task designed to probe sequential, multi-goal path generation based on experience acquired through unguided exploration.

##### Environment and actions

The environment is a square grid of size 50 *×* 50, with integer-valued coordinates (*r, c*) *∈ {* 0, …, 49 *}*^2^. Each grid cell corresponds to a unique state. At every time step, the agent deterministically selects one of four actions: *up, down, left*, or *right*, resulting in a transition to one of the four von Neumann neighboring cells. If an action would move the agent outside the grid, the position is reflected at the boundary, ensuring that all transitions remain within the environment.

##### Exploration phase

During training, the agent explores the environment through unguided random walks. We sample 10^5^ exploration episodes, each starting from a uniformly random grid location and consisting of a trajectory of fixed length (500 steps). At each step, one of the four actions is chosen uniformly at random.

For each episode, we record the set of grid locations visited at least once during the trajectory. Temporal order and visitation frequency within an episode are not retained; instead, each episode is represented as a binary indicator over grid locations, or equivalently an OR over grid locations. The learning process is as in the standard ECM algorithm.

##### Goal configuration

After exploration, the agent is evaluated on a foraging task. The starting location is fixed at the center of the grid, (24, 24). Four target locations are placed at equal Euclidean distance from the start and arranged approximately on a circle surrounding it, resulting in a symmetric configuration in which all target points are equidistant from the initial state. The behavioral target (corresponding to the goal in the preceding demonstrations) is defined by the set of all EMs that visit any of the four target points. Once one of these target points is reached, the EMs in which this point had been visited are removed (or equivalently, the corresponding index neurons are inhibited).

##### Utility definition

Utilities are defined as in the case of a single goal point, but for a different set of EMs. In this case the total state embedding is defined as all index cells that contains at least one of the target points that have not yet been reached. The same **W** matrix is used that maps the state embedding to the action utility. Target points that have not yet been reached are termed *active*.

##### Greedy action selection and sequential target removal

Starting from the current location, the agent constructs a path using a greedy action-selection policy. At each step, the four neighboring states are evaluated and the agent selects the action leading to the state with the highest utility. Ties are broken randomly. This process continues until an active target is reached or a maximum step limit is exceeded.

## References

Achiam, J., D. Held, A. Tamar, and P. Abbeel 2017. Constrained policy optimization. In International conference on machine learning, pp. 22–31. Pmlr.

Adcock, R.A., A. Thangavel, S. Whitfield-Gabrieli, B. Knutson, and J.D. Gabrieli. 2006. Reward-motivated learning: mesolimbic activation precedes memory formation. Neuron 50 (3): 507–517.

Andrychowicz, M., F. Wolski, A. Ray, J. Schneider, R. Fong, P. Welinder, B. McGrew, J. Tobin, O. Pieter Abbeel, and W. Zaremba. 2017. Hindsight experience replay. Advances in neural information processing systems 30.

Barreto, A., W. Dabney, R. Munos, J.J. Hunt, T. Schaul, H. Van Hasselt, and D. Silver. 2017. Successor features for transfer in reinforcement learning. Advances in neural information processing systems 30.

Bartol Jr, T.M., C. Bromer, J. Kinney, M.A. Chirillo, J.N. Bourne, K.M. Harris, and T.J. Sejnowski. 2015. Nanoconnectomic upper bound on the variability of synaptic plasticity. elife 4: e10778.

Behrens, T.E., T.H. Muller, J.C. Whittington, S. Mark, A.B. Baram, K.L. Stachenfeld, and Z. Kurth-Nelson. 2018. What is a cognitive map? organizing knowledge for flexible behavior. Neuron 100 (2): 490–509.

Bethus, I., D. Tse, and R.G. Morris. 2010. Dopamine and memory: modulation of the persistence of memory for novel hippocampal nmda receptor-dependent paired associates. Journal of Neuroscience 30 (5): 1610–1618.

Bhatnagar, S., R.S. Sutton, M. Ghavamzadeh, and M. Lee. 2009. Natural actor–critic algorithms. Automatica 45 (11): 2471–2482.

Biderman, N., A. Bakkour, and D. Shohamy. 2020. What are memories for? the hippocampus bridges past experience with future decisions. Trends in Cognitive Sciences 24 (7): 542–556.

Bittner, K.C., A.D. Milstein, C. Grienberger, S. Romani, and J.C. Magee. 2017. Behavioral time scale synaptic plasticity underlies ca1 place fields. Science 357 (6355): 1033–1036.

Blundell, C., B. Uria, A. Pritzel, Y. Li, A. Ruderman, J.Z. Leibo, J. Rae, D. Wierstra, and D. Hassabis. 2016. Model-free episodic control. arXiv preprint arXiv:1606.04460.

Bonini, L., C. Rotunno, E. Arcuri, and V. Gallese. 2022. Mirror neurons 30 years later: implications and applications. Trends in cognitive sciences 26 (9): 767–781.

Bornstein, A.M., M.W. Khaw, D. Shohamy, and N.D. Daw. 2017. Reminders of past choices bias decisions for reward in humans. Nature Communications 8 (1): 15958.

Bornstein, A.M. and K.A. Norman. 2017. Reinstated episodic context guides sampling-based decisions for reward. Nature neuroscience 20 (7): 997–1003.

Boscheron, J., P. Schoenmakers, A. Trivier, F. Lance, B. Herbelin, D. Van De Ville, M. Babo-Rebelo, and O. Blanke. 2026. Re-membering: spontaneous reactivation of motor cortex during memory re-experiencing. bioRxiv preprint.

Chen, L., K. Lu, A. Rajeswaran, K. Lee, A. Grover, M. Laskin, P. Abbeel, A. Srinivas, and I. Mordatch. 2021. Decision transformer: Reinforcement learning via sequence modeling. Advances in neural information processing systems 34: 15084–15097.

Chen, Z.S. and M.A. Wilson. 2023. How our understanding of memory replay evolves. Journal of Neurophysiology 129 (3): 552–580.

Chettih, S.N., E.L. Mackevicius, S. Hale, and D. Aronov. 2024. Barcoding of episodic memories in the hippocampus of a food-caching bird. Cell 187 (8): 1922–1935.

Christiano, P.F., J. Leike, T. Brown, M. Martic, S. Legg, and D. Amodei. 2017. Deep reinforcement learning from human preferences. Advances in neural information processing systems 30.

Costa, K.M. and G. Schoenbaum. 2025. Dopamine beyond temporal-difference reinforcement learning, Handbook of Behavioral Neuroscience, Volume 32, 251–261. Elsevier.

Dayan, P. 1993. Improving generalization for temporal difference learning: The successor representation. Neural computation 5 (4): 613–624.

Eichenbaum, H. 2017. Memory: organization and control. Annual review of psychology 68: 19–45.

Finn, C., P. Abbeel, and S. Levine 2017. Model-agnostic meta-learning for fast adaptation of deep networks. In International conference on machine learning, pp. 1126–1135. PMLR.

Forestier, S., R. Portelas, Y. Mollard, and P.Y. Oudeyer. 2022. Intrinsically motivated goal exploration processes with automatic curriculum learning. Journal of Machine Learning Research 23 (152): 1–41.

Freire, I.T., A.F. Amil, and P.F. Verschure. 2025. Sequential memory improves sample and memory efficiency in episodic control. Nature Machine Intelligence 7 (1): 43–55.

Frenkel, C., M. Lefebvre, J.D. Legat, and D. Bol. 2018. A 0.086-mm2 12.7-pj/sop 64k-synapse 256-neuron online-learning digital spiking neuromorphic processor in 28-nm cmos. IEEE transactions on biomedical circuits and systems 13 (1): 145–158.

Fried, I. 2022. Neurons as will and representation. Nature Reviews Neuroscience 23 (2): 104–114.

Gelbard-Sagiv, H., R. Mukamel, M. Harel, R. Malach, and I. Fried. 2008. Internally generated reactivation of single neurons in human hippocampus during free recall. Science 322 (5898): 96–101.

Gershman, S.J., J.A. Assad, S.R. Datta, S.W. Linderman, B.L. Sabatini, N. Uchida, and L. Wilbrecht. 2024. Explaining dopamine through prediction errors and beyond. Nature neuroscience 27 (9): 1645–1655.

Gershman, S.J. and N.D. Daw. 2017. Reinforcement learning and episodic memory in humans and animals: an integrative framework. Annual review of psychology 68: 101–128.

Greenstreet, F., H.M. Vergara, Y. Johansson, S. Pati, L. Schwarz, S.C. Lenzi, J.P. Geerts, M. Wisdom, A. Gubanova, L.B. Rollik, et al. 2025. Dopaminergic action prediction errors serve as a value-free teaching signal. Nature 643 (8074): 1333–1342.

Hu, H., J. Ye, G. Zhu, Z. Ren, and C. Zhang 2021. Generalizable episodic memory for deep reinforcement learning. In M. Meila and T. Zhang (Eds.), Proceedings of the 38th International Conference on Machine Learning, Volume 139, pp. 4380–4390.

Jeong, H., A. Taylor, J.R. Floeder, M. Lohmann, S. Mihalas, B. Wu, M. Zhou, D.A. Burke, and V.M.K. Namboodiri. 2022. Mesolimbic dopamine release conveys causal associations. Science 378 (6626): eabq6740.

John, T., Y. Zhou, A. Aljishi, B. Rieck, N.B. Turk-Browne, and E.C. Damisah. 2025. Representation of visual sequences in the tuning and topology of neuronal activity in the human hippocampus. bioRxiv preprint.

Jolliffe, I. 2011. Principal component analysis, International encyclopedia of statistical science, 1094–1096. Springer.

Kaelbling, L.P. 1993. Learning to achieve goals. In Ijcai, Volume 2, pp. 1094–8.

Kim, H., M.R. Mahmoodi, H. Nili, and D.B. Strukov. 2021. 4k-memristor analog-grade passive crossbar circuit. Nature communications 12 (1): 5198.

Kim, J., A. Joshi, L. Frank, and K. Ganguly. 2023. Cortical–hippocampal coupling during manifold exploration in motor cortex. Nature 613 (7942): 103–110.

Klaus, A., J. Alves da Silva, and R.M. Costa. 2019. What, if, and when to move: basal ganglia circuits and self-paced action initiation. Annual review of neuroscience 42 (1): 459–483.

Kolibius, L.D., F. Roux, G. Parish, M. Ter Wal, M. Van Der Plas, R. Chelvarajah, V. Sawlani, D.T. Rollings, J.D. Lang, S. Gollwitzer, et al. 2023. Hippocampal neurons code individual episodic memories in humans. Nature human behaviour 7 (11): 1968–1979.

Laskin, M., L. Wang, J. Oh, E. Parisotto, S. Spencer, R. Steigerwald, D. Strouse, S. Hansen, A. Filos, E. Brooks, et al. 2022. In-context reinforcement learning with algorithm distillation. arXiv preprint arXiv:2210.14215.

Lee, R.S., Y. Sagiv, B. Engelhard, I.B. Witten, and N.D. Daw. 2024. A feature-specific prediction error model explains dopaminergic heterogeneity. Nature neuroscience 27 (8): 1574–1586.

Lenci, A. 2018. Distributional models of word meaning. Annual review of Linguistics 4: 151–171.

Lengyel, M. and P. Dayan. 2007. Hippocampal contributions to control: the third way. Advances in neural information processing systems 20.

Li, C., M. Hu, Y. Li, H. Jiang, N. Ge, E. Montgomery, J. Zhang, W. Song, N. Dávila, C.E. Graves, et al. 2017. Analogue signal and image processing with large memristor crossbars. Nature electronics 1 (1): 52–59.

Li, Z., D. Zhu, Y. Hu, X. Xie, L. Ma, Y. Zheng, Y. Song, Y. Chen, and J. Zhao 2023. Neural episodic control with state abstraction. In The Eleventh International Conference on Learning Representations.

Lin, H., Y. Yang, R. Zhao, G. Pezzulo, and W. Maass. 2026. Neural sampling from cognitive maps enables goal-directed imagination and planning. Nature Machine Intelligence: in press.

Liu, G., M. Tang, and B. Eysenbach. 2024. A single goal is all you need: Skills and exploration emerge from contrastive rl without rewards, demonstrations, or subgoals. arXiv preprint arXiv:2408.05804.

Mackin, C., M.J. Rasch, A. Chen, J. Timcheck, R.L. Bruce, N. Li, P. Narayanan, S. Ambrogio, M. Le Gallo, S. Nandakumar, et al. 2022. Optimised weight programming for analogue memory-based deep neural networks. Nature communications 13 (1): 3765.

Markowitz, J.E., W.F. Gillis, M. Jay, J. Wood, R.W. Harris, R. Cieszkowski, R. Scott, D. Brann, D. Koveal, T. Kula, et al. 2023. Spontaneous behaviour is structured by reinforcement without explicit reward. Nature 614 (7946): 108–117.

Michelmann, S., B.P. Staresina, H. Bowman, and S. Hanslmayr. 2019. Speed of time-compressed forward replay flexibly changes in human episodic memory. Nature human behaviour 3 (2): 143–154.

Milani, S., N. Topin, M. Veloso, and F. Fang. 2024. Explainable reinforcement learning: A survey and comparative review. ACM Computing Surveys 56 (7): 1–36.

Milstein, A.D., Y. Li, K.C. Bittner, C. Grienberger, I. Soltesz, J.C. Magee, and S. Romani. 2021. Bidirectional synaptic plasticity rapidly modifies hippocampal representations. Elife 10: e73046.

Mukamel, R., A.D. Ekstrom, J. Kaplan, M. Iacoboni, and I. Fried. 2010. Single-neuron responses in humans during execution and observation of actions. Current biology 20 (8): 750–756.

Nicholas, J. and M.G. Mattar. 2026. Episodic memory facilitates flexible decision-making via access to detailed events. Nature Human Behaviour 10 (4): 760–776.

Piray, P. and N.D. Daw. 2021. Linear reinforcement learning in planning, grid fields, and cognitive control. Nature communications 12 (1): 4942.

Poh, J.H., M.A.T. Vu, J.K. Stanek, A. Hsiung, T. Egner, and R.A. Adcock. 2022. Hippocampal convergence during anticipatory midbrain activation promotes subsequent memory formation. Nature Communications 13 (1): 6729.

Pritzel, A., B. Uria, S. Srinivasan, A.P. Badia, O. Vinyals, D. Hassabis, D. Wierstra, and C. Blundell 2017. Neural episodic control. In International Conference on Machine Learning, pp. 2827–2836. PMLR.

Purandare, C. and M. Mehta. 2023. Mega-scale movie-fields in the mouse visuo-hippocampal network. Elife 12: RP85069.

Ramot, A., F.H. Taschbach, Y.C. Yang, Y. Hu, Q. Chen, B.C. Morales, X.C. Wang, A. Wu, K.M. Tye, M.K. Benna, et al. 2025. Motor learning refines thalamic influence on motor cortex. Nature 643 (8072): 725–734.

Robinson, N.T., L.A. Descamps, L.E. Russell, M.O. Buchholz, B.A. Bicknell, G.K. Antonov, J.Y. Lau, R. Nutbrown, C. Schmidt-Hieber, and M. Häusser. 2020. Targeted activation of hippocampal place cells drives memory-guided spatial behavior. Cell 183 (6): 1586–1599.

Rosman, B., M. Hawasly, and S. Ramamoorthy. 2016. Bayesian policy reuse. Machine Learning 104 (1): 99–127.

Roth, R.H. and J.B. Ding. 2024. Cortico-basal ganglia plasticity in motor learning. Neuron 112 (15): 2486–2502.

Russell, S. and P. Norvig. 2020. Artificial intelligence: A modern approach, 4th Edition. Pearson.

Samavat, M., T.M. Bartol, K.M. Harris, and T.J. Sejnowski. 2024. Synaptic information storage capacity measured with information theory. Neural Computation 36 (5): 781–802.

Schaul, T., D. Horgan, K. Gregor, and D. Silver 2015. Universal value function approximators. In International conference on machine learning, pp. 1312–1320. PMLR.

Sharpe, M.J., H.M. Batchelor, L.E. Mueller, C. Yun Chang, E.J. Maes, Y. Niv, and G. Schoenbaum. 2020. Dopamine transients do not act as model-free prediction errors during associative learning. Nature communications 11 (1): 106.

Shohamy, D. and R.A. Adcock. 2010. Dopamine and adaptive memory. Trends in cognitive sciences 14 (10): 464–472.

Song, W., M. Rao, Y. Li, C. Li, Y. Zhuo, F. Cai, M. Wu, W. Yin, Z. Li, Q. Wei, et al. 2024. Programming memristor arrays with arbitrarily high precision for analog computing. Science 383 (6685): 903–910.

Steunebrink, B.R., K.R. Thórisson, and J. Schmidhuber 2016. Growing recursive self-improvers. In International Conference on Artificial General Intelligence, pp. 129–139. Springer.

Stickgold, R. and M.P. Walker. 2013. Sleep-dependent memory triage: evolving generalization through selective processing. Nature neuroscience 16 (2): 139–145.

Stöckl, C., Y. Yang, and W. Maass. 2024. Local prediction-learning in high-dimensional spaces enables neural networks to plan. Nature Communications 15 (1): 2344.

Sun, W., J. Winnubst, M. Natrajan, C. Lai, K. Kajikawa, A. Bast, M. Michaelos, R. Gattoni, C. Stringer, D. Flickinger, et al. 2025. Learning produces an orthogonalized state machine in the hippocampus. Nature 640 (8057): 165–175.

Sutton, R.S. and A.G. Barto. 2018. Reinforcement learning: An introduction. MIT press.

Sze, V., Y.H. Chen, T.J. Yang, and J.S. Emer. 2017. Efficient processing of deep neural networks: A tutorial and survey. Proceedings of the IEEE 105 (12): 2295–2329.

Sze, V., Y.H. Chen, T.J. Yang, and J.S. Emer. 2020. How to evaluate deep neural network processors: Tops/w (alone) considered harmful. IEEE Solid-State Circuits Magazine 12 (3): 28–41.

Tacikowski, P., G. Kalender, D. Ciliberti, and I. Fried. 2024. Human hippocampal and entorhinal neurons encode the temporal structure of experience. Nature 635 (8037): 160–167.

Taipalus, T. 2024. Vector database management systems: Fundamental concepts, use-cases, and current challenges. Cognitive Systems Research 85: 101216.

Takahashi, S. 2015. Episodic-like memory trace in awake replay of hippocampal place cell activity sequences. Elife 4: e08105.

Teyler, T.J. and P. DiScenna. 1986. The hippocampal memory indexing theory. Behavioral neuroscience 100 (2): 147.

Vellani, V., L.P. de Vries, A. Gaule, and T. Sharot. 2020. A selective effect of dopamine on information-seeking. Elife 9: e59152.

Waydo, S., A. Kraskov, R.Q. Quiroga, I. Fried, and C. Koch. 2006. Sparse representation in the human medial temporal lobe. Journal of Neuroscience 26 (40): 10232–10234.

Wittgenstein, L. 2009. Philosophical Investigations (Revised). Wiley-Blackwell.

Wu, Y. and W. Maass. 2025. A simple model for behavioral time scale synaptic plasticity (btsp) provides content addressable memory with binary synapses and one-shot learning. Nature communications 16 (1): 342.

Xiao, K., Y. Li, B.J. Sullivan, G. Li, and J.C. Magee. 2025. Rapid neocortical network modifications via dendritic plateau potential induced plasticity. bioRxiv preprint.

Yaeger, C.E., R. Mojica Soto-Albors, W. Liu, A. Beltramini, and M.T. Harnett. 2025. Plateau potentials are instructive signals for behavioral timescale synaptic plasticity in the neocortex. bioRxiv preprint.

Yan, A. and Z. Cheng. 2024. A review of the development and future challenges of case-based reasoning. Applied Sciences 14 (16): 7130.

Yang, Y. and W. Maass. 2026. The inherent capacity of neurons to learn order relations and support abstract reasoning. Nature Communications 17: 9436. 10.1038/s41467-026-76102-5.

Zhou, P., A.J. Edwards, F.B. Mancoff, S. Aggarwal, S.K. Heinrich-Barna, and J.S. Friedman. 2025. Neuromorphic hebbian learning with magnetic tunnel junction synapses. Communications Engineering 4 (1): 142.

Zintgraf, L., K. Shiarlis, M. Igl, S. Schulze, Y. Gal, K. Hofmann, and S. Whiteson. 2019. Varibad: A very good method for bayes-adaptive deep rl via meta-learning. arXiv preprint arXiv:1910.08348.

Zutshi, I., A. Apostolelli, W. Yang, Z.S. Zheng, T. Dohi, E. Balzani, A.H. Williams, C. Savin, and G. Buzsáki. 2025. Hippocampal neuronal activity is aligned with action plans. Nature 639: 153–161. 10.1038/s41586-024-08397-7.

